# Continuous Directed Evolution of a Plant Histidinol Dehydrogenase to Extend Lifespan

**DOI:** 10.1101/2025.07.01.662441

**Authors:** Edmar R. Oliveira-Filho, Anuran K. Gayen, Bryan J. Leong, Katharina Belt, A. Harvey Millar, Andrew D. Hanson

## Abstract

Enzyme protein turnover accounts for about half the maintenance energy budget in plants. Slowing turnover – i.e., extending lifespan – of short-lived enzymes is thus a rational strategy to conserve energy and carbon, and raise crop productivity. *Arabidopsis* histidinol dehydrogenase (HDH) is a short-lived enzyme that can sustain life-shortening damage from its aminoaldehyde reaction intermediate. We used the yeast OrthoRep continuous directed evolution system in a *his4*Δ strain to raise HDH protein abundance (a proxy for lifespan) by selecting for growth rate while tapering histidinol concentration and escalating that of the inhibitor histamine. Improved HDHs carried diverse nonsynonymous mutations and ranged 20-fold in level. Improved HDH performance was associated with higher HDH abundance in some cases and with greater catalytic efficiency or histamine resistance in others. These findings indicate that OrthoRep-based directed evolution can extend enzyme lifespan *in vivo* in addition to, as expected, altering kinetic properties.

## INTRODUCTION

The continuous degradation and resynthesis (i.e., turnover) of enzyme proteins is a major item in the energy budget of all organisms^1,2^ and in plants this enzyme replacement can represent up to 50% of maintenance energy costs.^3^ If these costs could be cut by extending enzyme lifespan, the energy – and hence carbon – saved could be used to support growth and yield, i.e., to boost crop productivity.^4,5^ A useful metric for enzyme lifespan is the ‘Catalytic Cycles till Replacement’ (CCR) value, which is the number of catalytic reactions that an enzyme mediates during its lifetime (as estimated from metabolic flux rate/enzyme replacement rate).^2,6^ The median CCR for *Arabidopsis* enzymes is 4 × 10^5^ but many have CCRs <4 × 10^4^.^6^ These short-lived enzymes typically have chemically reactive substrates, products, or intermediates and are thus prone to self-inactivate during catalysis due to chemical attack on vulnerable residues in or near the active site.^6,7^ Enzymes like this are good candidates for life-extension because they can potentially be hardened against inactivation, e.g., by replacing vulnerable residues with resistant ones,^7^ and are even better candidates if they are abundant as energy saved by longer life scales with amount of protein present.^1,8^

*Arabidopsis* histidinol dehydrogenase (HDH, EC 1.1.1.23) is one such enzyme. HDH mediates the last two steps in histidine biosynthesis, the NAD^+^-dependent oxidation of histidinol to histidine via histidinal (**Figure 1A**).^9^ HDH has a low CCR (3.3 × 10^4^),^6^ is in the topmost protein abundance decile,^10^ and has a reactive aminoaldehyde intermediate^11^ that attacks HDH protein residues.^12,13^ We therefore sought to extend HDH lifespan via continuous directed evolution in the yeast (*Saccharomyces cerevisiae*) OrthoRep system^14,15^ using abundance as a proxy for lifespan. The logic is this.^5^ (i) Protein abundance is set by rates of transcription, translation, and degradation. (ii) Major increases in transcription and translation rates of the target gene are unlikely.^15^ (iii) Slower degradation rate, i.e., longer lifespan, is therefore predicted to have a substantial role in increasing abundance.

**Figure 1.**
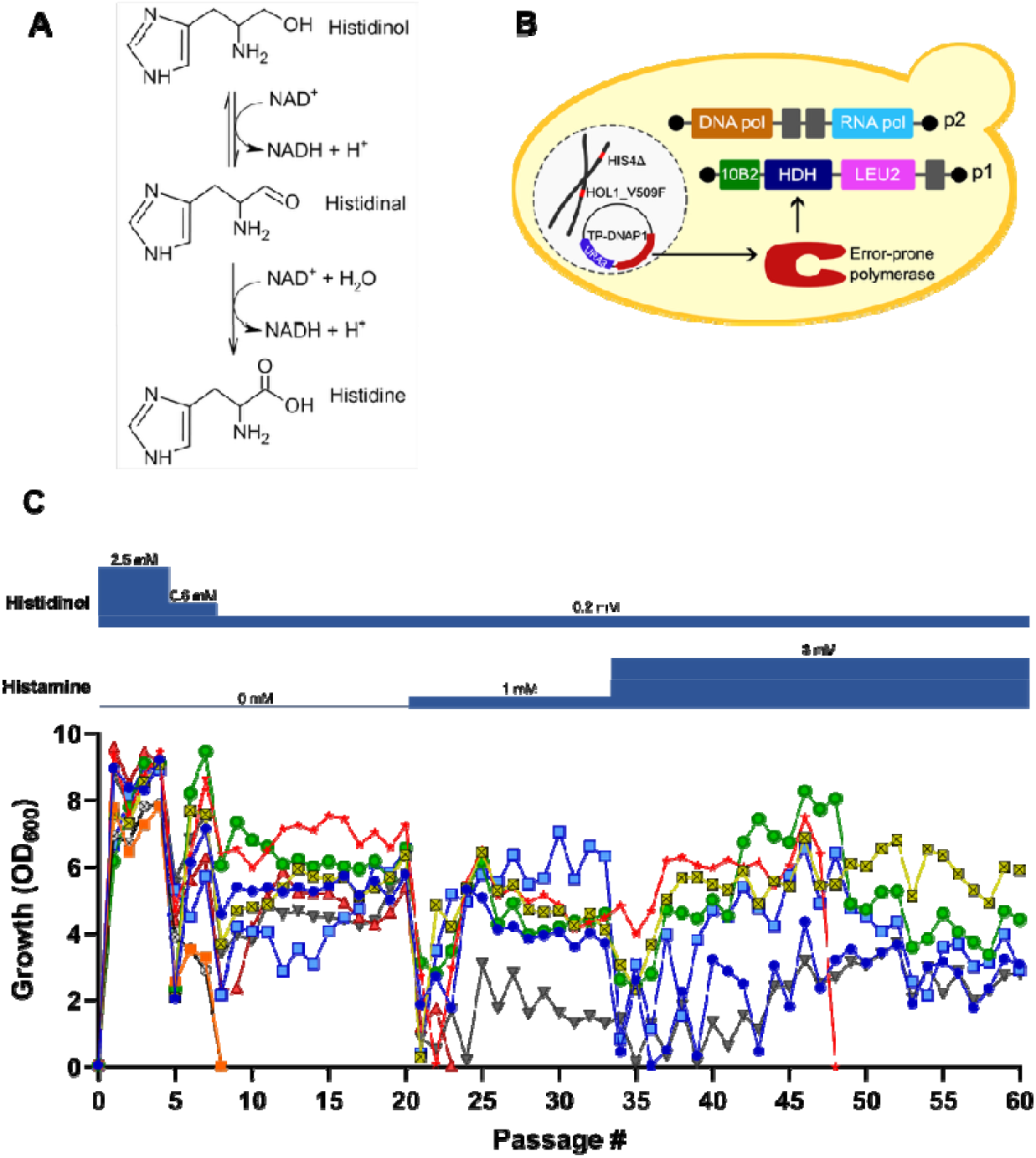
Continuous directed evolution of HDH in the OrthoRep system. (A) The two-step HDH reaction and the structures of histidinol, the aminoaldehyde intermediate, and the histidine product. (B) Overview of OrthoRep. The p1 plasmid carries the HDH target gene, expressed from the 10B2 promoter, and a *LEU2* marker. (C) Full campaign growth profiles for representative subpopulations (in different colors) expressing the polyadenylated construct. Cells received stepwise decreasing levels of histidinol, and stepwise increasing levels of histamine as indicated. Passage length was generally 2-3 days but was extended to 4-5 days for the early passages in each step except the first. Note that not all subpopulations survived as selection proceeded. Growth of representative subpopulations expressing the non-polyadenylated construct is shown in **Figure S1**.

OrthoRep (**Figure 1B**) offers mutational depth, scale, and durability, makes no off-target mutations, and is suitable for plant enzymes.^16^ The target enzyme is expressed from a linear cytoplasmic plasmid (p1) that is replicated by a p1-specific hypermutator DNA polymerase encoded by a nuclear plasmid; genes on p1 are transcribed by an RNA polymerase encoded by a second cytoplasmic plasmid (p2). Target enzyme activity is coupled to growth by using a platform strain in which the corresponding enzyme is knocked out (here, *his4*Δ), and evolution campaigns are run by selecting populations for growth rate.^14-16^ Selecting for target enzyme activity in this way is expected to lead not only to higher enzyme abundance but also to improved kinetic properties. However, for a primary metabolic enzyme acting on its normal substrate, such kinetic improvement is likely to be modest,^17,18^ leaving scope to select for longer life.^5^ The combined assessment of protein abundances and kinetic properties allowed dissection of these contributions for different mutants.

## RESULTS AND DISCUSSION

### Platform Strain Construction and Check on Complementation by HDH

Yeast cannot take up histidinol unless the Hol1 transporter carries a mutation such as V509F.^19^ Homologous recombination was therefore used to introduce the *HOL1*-V509F mutation into a *his4*Δ knockout of strain BY4742 harboring HDH (recoded for yeast, minus the plastid targeting peptide) in the CEN6/ARS4 nuclear plasmid. The resulting strain grew when given 2.5 mM histidinol. (Exogenous histidinol is the only source of histidinol in a *his4*Δ knockout because His4 is a trifunctional histidine biosynthesis protein that carries the upstream enzymes phosphoribosyl-ATP pyrophosphatase and phosphoribosyl-AMP cyclohydrolase as well as HDH.) The plasmid was then purged prior to installing the OrthoRep system (see below). The CEN6/ARS4 plasmid’s TDH3 promoter drives expression at least twice that of the p1 plasmid’s 10B2 promoter;^20^ target genes must pass this nuclear plasmid complementation check for the OrthoRep system to be likely to work.^16^

### Directed Evolution Campaigns

HDH, preceded by the 10B2 promoter and plus or minus a 72A poly(A) tail (to give higher and lower expression levels, respectively^16^) was introduced into the p1 plasmid by homologous recombination in the p1 donor strain GA-Y319.^21^ Recombinant p1 plasmids and wildtype p2 plasmid were then transferred by protoplast fusion from GA-Y319 to the above BY4742 *HOL1*-V509F *his4*Δ strain harboring error-prone DNA polymerase TP-DNAP1.^16^ For the constructs with or without a poly(A) tail, 9 independent populations from separate protoplast fusion events were used to launch evolution campaigns in medium containing 2.5 mM histidinol. Of the 18 initial populations, 15 reached an OD_600_ of 7–10 by the end of each passage (**Figure 1C** and **Figure S1**). Selection was intensified for these 15 populations, first by reducing histidinol concentration stepwise to 0.2 mM, then by increasing histamine concentration stepwise from zero to 3 mM. Histamine is a potent competitive inhibitor of HDH;^22^ selecting for resistance to such inhibitory analogs is a classical strategy to raise enzyme abundance or activity.^23,24^ As selection intensified, growth of surviving cultures improved during successive passages (**Figure 1C** and **Figure S1**). Five populations with the poly(A) tail and two populations without it survived until campaigns ended (after 60 passages). Most of these populations were split into subpopulations along the way to ensure some redundancy and to let successful populations evolve independently after the split.

### Campaign Outcomes

Bulk sequencing of cultures at the end of campaigns detected 23 nonsynonymous mutations that had swept subpopulations, i.e., fully replaced a wildtype base and its cognate residue (**Figure 2A**). None of these mutations recurred in independent populations, i.e., there were no cases of convergent evolution. Three mutant HDHs carried single nonsynonymous mutations while 18 carried two or more (**Figure 2A**). As expected, these mutations were all transitions in the first or second base of the codon.^14,25^ Occasional mutations in the 10B2 promoter (**Figure 2A**) and some synonymous mutations (**Figure S2**) were also found.

**Figure 2.**
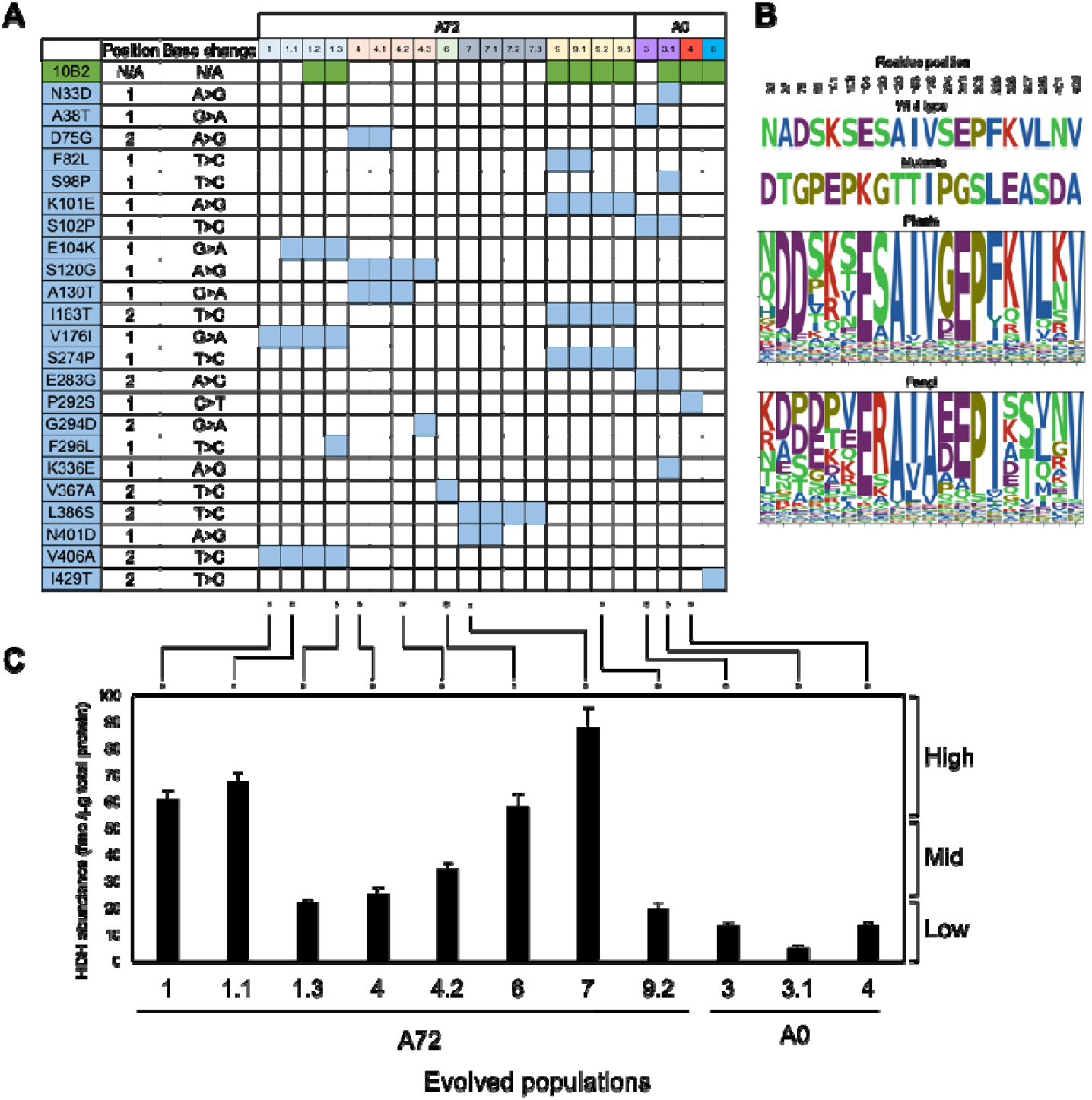
Nonsynonymous mutations found in evolved populations expressing the polyadenylated (A72) or non-polyadenylated (A0) construct. (A) The mutations and the subpopulations they came from. (B) Wildtype and mutant residues compared to sequence logos for plant and fungal HDHs based on 100 sequences from diverse taxa in each kingdom. (C) Abundance at the end of campaigns of the HDH protein in subpopulations harboring various mutant HDHs. The classification into high-, low-, and mid-abundance is shown on the right. Data for both nonsynonymous and synonymous mutations are in **Figure S2**.

To evaluate whether the observed mutations occur in nature, we compared the mutant HDHs with sequences representing the diversity in plant and fungal HDHs, plotted as sequence logos (**Figure 2B**). Some mutations (e.g., S98P and K336E) are at variable positions and occur naturally, and so are likely to be neutral or nearly so. Others (e.g., A130T, V406A) are at highly conserved positions and absent in nature. Mutations like this (i.e., that lie outside natural sequence space but conserve enzyme function) may well specify significant improvements^26^ and so were analyzed further.

### HDH Abundance

Eleven mutant sequences with at least one non-natural mutation at a conserved position (**Figure S3**) were chosen for analysis of HDH abundance in subpopulations at the end of campaigns. Targeted multiple reaction monitoring of HDH derived-peptides by liquid chromatography-mass spectrometry showed a 20-fold range of HDH abundance across the populations, with the highest value (88 fmol μg^-1^ protein) equivalent to 0.4% of total protein, or fivefold more than the native His4 protein^10^ that HDH replaces (**Figure 2C**). The mutant sequences were classified as high-, low-, or mid-abundance (**Figure 2C**). An untargeted data-dependent proteomics analysis of the same samples showed that the yeast His4 protein was effectively absent (thereby confirming the *HIS4* knockout), and that there were no consistent changes in other histidine biosynthesis enzymes (**Figure S4**).

### HDH Enzyme Kinetics

Six evolved HDH sequences representing various abundances were expressed in *Escherichia coli* and compared with wildtype HDH with respect to *K*_M_ and *k*_cat_ values and to sensitivity to inhibition by histamine, scored as the concentration that inhibited activity by 50% (*i*_50_) (**Table 1** and **Figure 3A**). High-abundance HDH A72_7 had *K*_M_, *k*_cat_, and *i*_50_ values close to wildtype HDH, making abundance (i.e., lifespan) alone a sufficient explanation for superior *in vivo* performance in this case. The two other high-abundance HDHs had altered kinetics: A72_6 was strongly histamine-resistant (*i*_50_ >50-fold more than wildtype) and had a 5-fold higher *K*_M_ for histidinol; A72_1.1 was also histamine-resistant (*i*_50_ 10-fold more than wildtype). Mid-abundance A72_4 and low-abundance A0_3.1 also had altered kinetics; both had *K*_M_ values ∼4-fold lower than wildtype. In contrast, low-abundance A0_4 showed no kinetic differences from wildtype. As *S. cerevisiae* has some capacity to catabolize histamine,^27,28^ upregulation of this capacity by a rare nuclear gene mutation^14^ could in principle explain such an occasional outlier.

**Table 1.** Abundance and Kinetic Parameters of Wildtype and Evolved Arabidopsis HDHs^*ab*^.

| Enzyme | Abundance (fmol $\mu\text{g}^{-1}$) | Abundance class | $K_M$ ($\mu\text{M}$) | $k_{\text{cat}}$ ($\text{s}^{-1}$) | $k_{\text{cat}}/K_M$ ($\text{s}^{-1} \text{mM}^{-1}$) | Histamine $i_{50}$ (mM) |
| --- | --- | --- | --- | --- | --- | --- |
| Wildtype | -- | -- | $20.9 \pm 2.4$ | $2.7 \pm 0.1$ | $131 \pm 12$ | $0.2 \pm 0.03$ |
| A72_1.1 | $67.5 \pm 3.4$ | High | $13.7 \pm 2.2$ | $1.6 \pm 0.1$ | $118 \pm 16$ | $1.8 \pm 0.15$ |
| A72_4 | $25.6 \pm 2.2$ | Mid | $5.9 \pm 1.5$ | $1.00 \pm 0.1$ | $172 \pm 58$ | $0.2 \pm 0.07$ |
| A72_6 | $58.5 \pm 4.6$ | High | $104.5 \pm 6.6$ | $2.7 \pm 0.1$ | $26 \pm 1.4$ | >10 |
| A72_7 | $87.9 \pm 7.3$ | High | $17.4 \pm 2.4$ | $2.3 \pm 0.1$ | $135 \pm 16$ | $0.3 \pm 0.07$ |
| A0_3.1 | $5.3 \pm 0.7$ | Low | $4.8 \pm 0.5$ | $1.1 \pm 0.1$ | $229 \pm 37$ | $0.2 \pm 0.02$ |
| A0_4 | $13.8 \pm 1.1$ | Low | $17.8 \pm 4.4$ | $2.6 \pm 0.2$ | $153 \pm 31$ | $0.2 \pm 0.02$ |

<sup>a</sup> Data are means of 3 or 4 replicates ± standard deviation.
<sup>b</sup> Histamine $i_{50}$ is the histamine concentration required for 50% inhibition of the reaction rate measured at a histidinol concentration of 20 $\mu\text{M}$ (i.e., the $K_M$ value for wildtype HDH).

**Figure 3.**
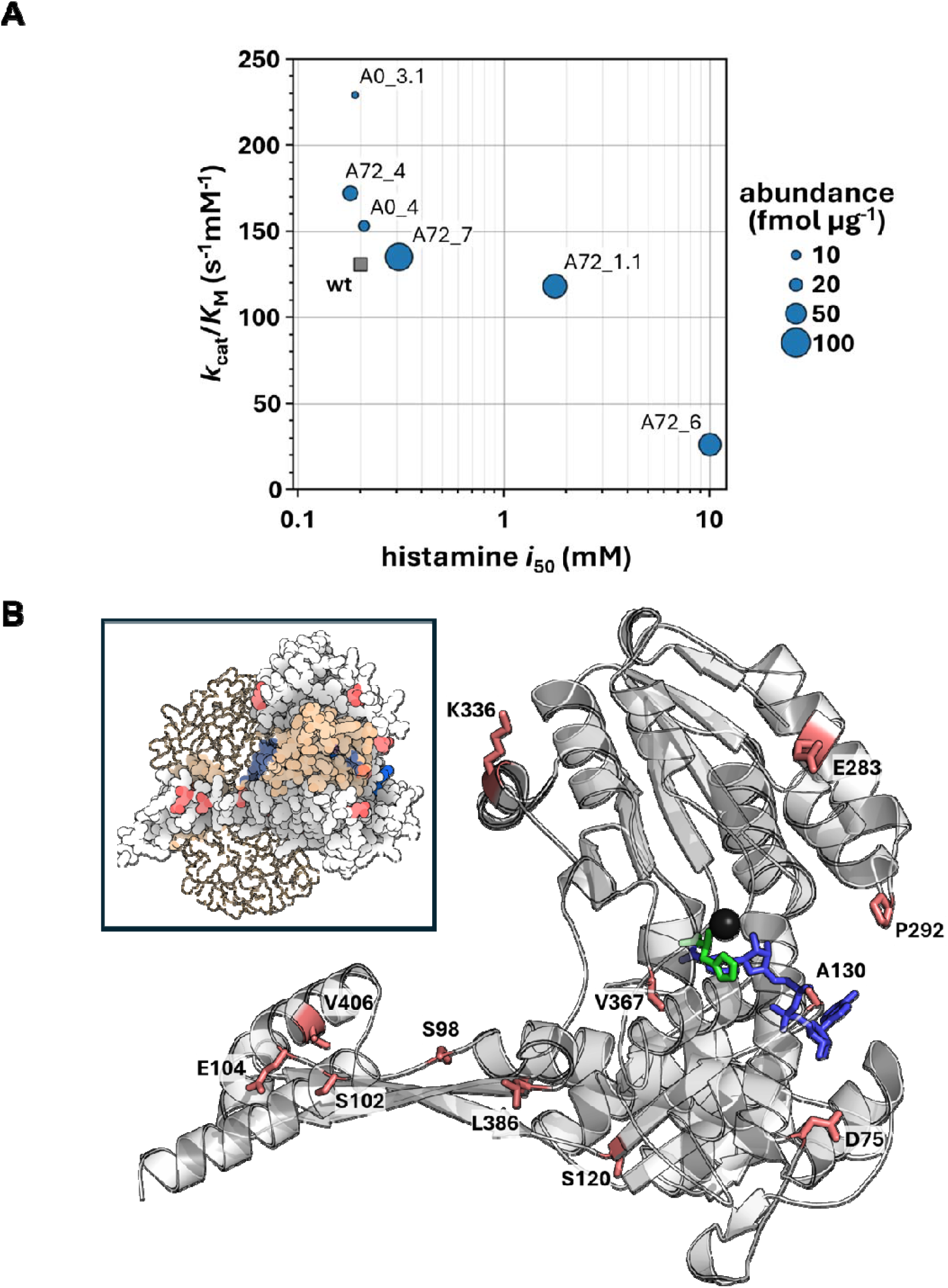
Evolutionary outcomes for representative HDH variants. (A) Scatter plot illustrating mutant enzyme kinetic properties, with histamine *i*_50_ on the *x*-axis and catalytic efficiency (*k*_cat_/*K*_M_) on the y-axis. Protein abundance is represented by the area of each circular symbol. Wild-type HDH is shown as a gray square. The *i*_50_ value for variant A72_6 is conservatively plotted at 10 mM. (B) Structural mapping of mutations onto the modeled *Arabidopsis* HDH monomer structure bound to Zn^2^+ (black) and NAD+ (blue). Mutated residues are highlighted in red, and the apparent position of L-histidine is shown in green. Inset: Space-filling representation of the HDH dimer, with monomers colored beige and white.

### Mutation Positions and Types

Plant HDHs are homodimers in which each monomer contains four domains; NAD^+^ binds to residues in domain I, while Zn^2+^ is coordinated in and histidinol binds to residues in domains II and IV.^9^ Mapping the positions of mutations in the 11 HDHs chosen for analysis of abundance (**Figure 3B**) onto the modeled structure of the *Arabidopsis* HDH monomer shows that all except the V367A mutation (which increased histamine resistance >50-fold) are near the protein surface, i.e., far from the active site and hence hard to rationalize individually in the absence of experimental structures for the mutant enzymes.^29^ Of note, however: the ratio of conservative to non-conservative mutations is 7:1 for the 4 most-abundant HDHs *vs*. 1:2 for the 4 least-abundant; this difference is statistically significant (χ^2^ test, *P* <0.01). Abundant HDHs thus carry fewer mutations that are liable to disrupt catalytic function or to destabilize structure and reduce lifespan and abundance.^30^

## CONCLUSIONS

This work explored whether the yeast OrthoRep continuous directed evolution system can increase the *in-vivo* lifespan of a short-lived plant enzyme. As measuring lifespan is challenging and costly,^2^ we used enzyme abundance as a proxy to screen for longer lifespan. We obtained mutant HDHs with a wide range of abundance, and – as is typical in directed evolution^5,24,31^ – with kinetic property changes (**Figure 3A** and **Table 1**). A large property change in two cases was resistance to the histamine used in selection to attenuate HDH activity (the aim having been to favor evolution of a compensatory increase in HDH protein level). Other kinetic changes were on the relatively modest scale expected from selection of an already-efficient enzyme acting on its physiological substrate.^5^ As **Figure 3A** shows, the evolutionary outcomes are distributed throughout the functional space defined by abundance, catalytic efficiency, and histamine resistance, with each variable contributing to a different extent. We therefore conclude that combining OrthoRep campaigns with the type of analysis in **Figure 3A** is an effective way to obtain and screen evolved enzymes with high abundance and to separate them from those whose improved performance is due mainly or wholly to altered kinetics. Enzymes retained by such a screen, e.g., HDH A72_7, can then be advanced to technically demanding and costly stable isotope measurements^2^ of *in-vivo* lifespan in yeast or a plant host.

A core tenet of this work is that OrthoRep can improve an enzyme’s cellular activity by increasing its abundance, with longer lifespan being *a priori* the major contributing factor. Of the other possible factors, i.e., higher transcription or translation rate, transcription rate is essentially excluded since there were few promoter mutations and these were associated with low, not high, abundance (**Figure 2A,C**). Synonymous mutations (**Figure S2**) can potentially raise the translation rate of commercially codon-optimized sequences in yeast,^32^ but as such increases are typically no more than 3-fold^33,34^ they are unlikely to explain the observed 20-fold range of abundance (**Figure 2C**).

## MATERIALS AND METHODS

### Media, Strains, Plasmids, and Primers

*E. coli* was grown in LB medium plus 100 μg/mL kanamycin or 50 μg/mL carbenicillin. Yeast was grown aerobically in YPD or synthetic complete (SC) selection medium as described.^16,35^ *Arabidopsis* HDH (At5g63890) was truncated, codon-optimized for *E. coli* or yeast, and synthesized by GenScript (Piscataway, NJ) (**Table S1**). Cloning operations were as described.^16,35^ Kits and enzymes were from Thermo Fisher Scientific (Waltham, MA) unless otherwise stated. Primers (**Table S2**) were from Eurofins Genomics (Louisville, KY).

### OrthoRep Campaigns

BY4742 *his4*Δ (Y13437) from Euroscarf (Oberursel, Germany) was transformed with ArEc-TDH3 (nuclear plasmid, *URA3* marker) harboring recoded HDH. Homologous recombination was then used to introduce the *HOL1*-V509F mutation.^36^ Briefly, wildtype *HOL1* was replaced by a *LEU2* marker cassette (assembled in pAGT572 using Golden Gate cloning), which was then replaced by *HOL1*-V509F (generated by site-directed mutagenesis). Clones were selected in SC -His -Ura -Trp with 2.5 mM histidinol, and sequence-verified *HOL1*-V509F clones were passaged 5 times in SC medium plus uracil to purge the nuclear plasmid. Yeast transformation and protoplast fusion were performed as described^16,37^ to construct the platform strain. The obtained Y13437 BY4742 *his3*Δ1 *leu2*Δ0 *lys2*Δ0 *ura3*Δ0 *his4*Δ *HOL1*-V509F strain harboring the 633 error-prone TP-DNAP in ArEc-TDH3 (nuclear plasmid, *URA3* marker) and p1_HDH (*LEU2* marker) was selected in SC -His -Ura -Leu -Trp containing 2.5 mM histidinol. Independent populations were subjected to evolution campaigns in the same medium containing histidinol (2.5–0.25 mM) and histamine (0–3 mM).^22,27^ Three-mL cultures were started at OD_600_ = 0.05 in 24-well plates, and incubated at 30°C, 800 rpm, 80% humidity in an HT Multitron shaker (Infors AG, Switzerland). Growth was monitored at OD_600_. Populations were subcultured every 3-7 days for 60 passages. Genetic manipulations and HDH mutations were verified by Sanger sequencing of PCR products.

### Protein Quantification *In Vivo*

Protein abundance was assessed at the end of campaigns. (Abundance at the start of campaigns is uninformative because p1 is a multicopy plasmid. Early in a campaign, cells can carry both recombinant p1 and residual wildtype p1. Wildtype p1 enables higher p1 copy numbers and higher expression of the recombinant p1-encoded gene of interest.^15^ Wildtype p1 lacks a selection marker and is subsequently purged from the population.) Selected HDH final populations were grown in triplicate in 50 mL SC -His -Leu -Ura plus 0.25 mM histidinol. When OD_600_ reached 2.5–3.5, cultures were harvested by centrifugation (5000*g*, 5 min, 4°C), washed twice with ice-cold water, frozen in liquid N_2_, and freeze-dried overnight. Sample preparation and mass spectrometry analysis methods were similar to those used previously^38^ and are given in Proteomics Methods (Supplemental Information).

### HDH Assays

HDH mutant sequences were generated by site-directed mutagenesis using KLD Enzyme Mix (New England Biolabs, Ipswich, MA) or synthesized by GenScript (**Table S1**), cloned into pET28b(+) (Novagen, Inc.), sequence-verified, and introduced into *E. coli* BL21 (DE3). Recombinant *E. coli* cells were grown in LB medium plus kanamycin at 37°C, 250 rpm. When OD_600_ reached ∼0.5, cultures were cold-shocked in an ice bath for 30 min, IPTG was added (final concentration 100 μM), and incubation was continued overnight at 18°C, 180 rpm. Cells were harvested by centrifugation (5000*g*, 15 min, 4°C) and stored at -80°C. Cell pellets were sonicated in 50 mM HEPES-NaOH (pH 7.5), 300 mM NaCl, 5 mM dithiothreitol, and 200 μM phenylmethanesulfonyl fluoride. Soluble fractions were collected by centrifugation (25000*g*, 15 min), desalted on PD-10 columns (Cytiva Life Sciences, Sweden) into the above buffer minus phenylmethanesulfonyl fluoride, plus 10% (w/v) glycerol, and stored at -80°C. Protein was determined by Bradford assay, and protein profiles (10 μg per sample) were visualized by SDS-PAGE on 12.5% gels stained with GelCode Blue Safe Protein Stain (Thermo Fisher Scientific). The quantity of HDH in each sample (as percent of total protein) was estimated using ImageJ software;^39^ values were 4% to 20%. HDH activity in desalted extracts was measured spectrophometrically in a reaction mix containing 500 mM Bicine-NaOH (pH 8.5), 5 mM NAD^+^, 0.5 mM MnCl_2_, 5 μg/mL HDH, and 0.001–1 mM histidinol. The histamine concentration required for 50% inhibition of the reaction rate (*i*_50_) was measured at a histidinol concentration of 20 μM (the *K*_M_ value of wildtype HDH).

### Structure Modeling

The dimeric structure of wild-type *Arabidopsis* HDH was predicted using AlphaFold 3, based on amino acid sequence. The prediction included two NAD+ molecules and two Zn^2+,^ ions and was generated using a random seed without structural templates. To approximate the binding site of L-histidine, the modeled HDH structure was superimposed onto the ligand-bound crystal structure of L-histidinol dehydrogenase from *Medicago truncatula* (PDB ID: 5VLD). Structural visualizations were prepared using PyMOL (Schrödinger, LLC) and Protein Imager.^40^

## Supporting information

Proteomics Methods

## Supporting information

### Proteomics Methods

**Figure S1**. Full campaign growth profiles for representative subpopulations that expressed the non-polyadenylated HDH construct.

**Figure S2**. Figure S2. Full listing of nonsynonymous (mid-blue) and synonymous (light-blue) mutations that swept subpopulations.

**Figure S3**. Figure S3. The 11 mutant HDH sequences selected for proteomics and (boxed in red) for kinetic analyses.

**Figure S4**. Untargeted (Orbitrap) proteomics data on levels of histidine biosynthesis enzymes in subpopulations at the end of evolution campaigns.

**Table S1**. Protein and nucleotide sequences of the HDHs used in this study.

**Table S2**. Primers used in this study.

**Table S3**. Peptide sequences and mass spectrometry analysis parameters used for targeted selected reaction monitoring analysis of *Arabidopsis* HDH.

## Author contributions

A.D.H. and E.R.O.-F. devised the study. E.R.O.-F. ran directed evolution campaigns and analyzed enzyme kinetics. B.J.L. developed enzyme assays. K.B. and A.H.M. conducted proteomics analyses. A.K.G. built structure models. All authors analyzed data. A.D.H. and E.R.O.-F. wrote the manuscript.

## ACKNOWLEDGEMENTS

We thank M.A. Wilson and N. Smith for discussions, and F. Rahimi for assistance with generating sequence logos. This work was supported by USDA National Institute of Food and Agriculture Hatch project FLA-HOS-005796, and an Endowment from the C.V. Griffin, Sr. Foundation. The protein analysis was supported by the Australian Research Council (FL200100057 to A.H.M) and infrastructure through Bioplatforms Australia as part of the National Collaborative Research Infrastructure Strategy (NCRIS) in Australia. We thank Elke Ströher from WA Proteomics Facility at the University of Western Australia for support with mass spectrometry analysis.

## For Table of Contents Only

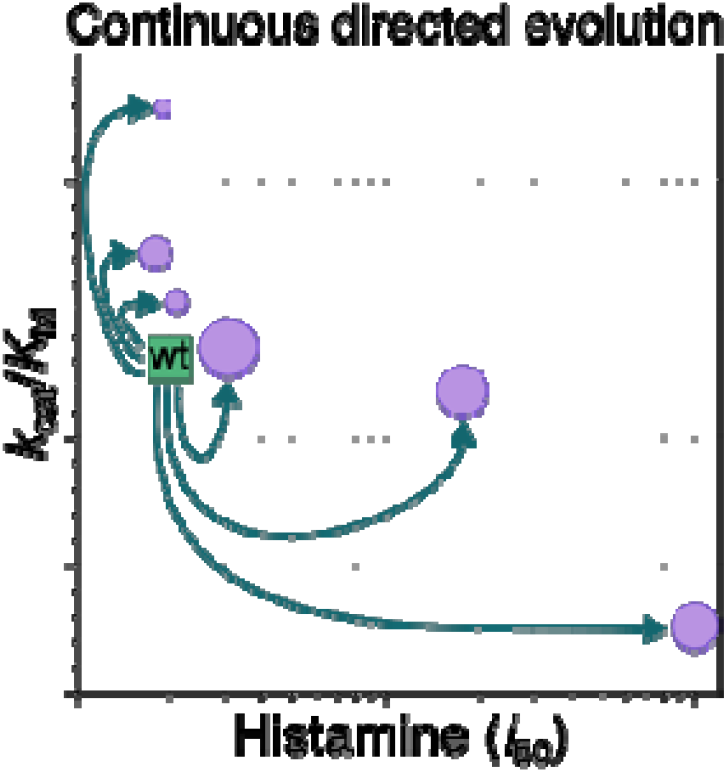

## REFERENCES

(1) Koehn, R. K. The cost of enzyme synthesis in the genetics of energy balance and physiological performance. Biol. J. Linn. Soc. Lond. 1991, 44, 231–247.

(2) Tivendale, N. D.; Hanson, A. D.; Henry, C. S.; Hegeman, A. D.; Millar, A. H. Enzymes as parts in need of replacement – and how to extend their working life. Trends Plant Sci. 2020, 25, 661–669.

(3) Amthor, J. S.; Bar-Even, A.; Hanson, A. D.; Millar, A. H.; Stitt, M.; Sweetlove, L. J.; Tyerman, S. D. Engineering strategies to boost crop productivity by cutting respiratory carbon loss. Plant Cell 2019, 31, 297–314.

(4) Reynolds, M.; Atkin, O. K.; Bennett, M.; Cooper, M.; Dodd, I. C.; Foulkes, M. J.; Frohberg, C.; Hammer, G.; Henderson, I. R.; Huang, B.; Korzun, V.; McCouch, S. R.; Messina, C. D.; Pogson, B. J.; Slafer, G. A.; Taylor, N. L.; Wittich, P. E. Addressing research bottlenecks to crop productivity. Trends Plant Sci. 2021, 26, 607–630.

(5) Joshi, J.; Amthor, J. S.; McCarty, D. R.; Messina, C. D.; Wilson, M. A., Millar, A. H.; Hanson A. D. Why cutting respiratory CO2 loss from crops is possible, practicable, and prudential. Mod. Agric. 2023, 1, 16–26.

(6) Hanson, A. D.; McCarty, D. R.; Henry, C. S.; Xian, X.; Joshi, J.; Patterson, J. A.; García-García, J. D.; Fleischmann, S. D.; Tivendale, N. D.; Millar, A. H. The number of catalytic cycles in an enzyme’s lifetime and why it matters to metabolic engineering. Proc. Natl. Acad. Sci. USA 2021, 118, e2023348118.

(7) Bathe, U.; Leong, B. J.; McCarty, D. R.; Henry, C. S.; Abraham, P. E.; Wilson, M. A.; Hanson, A. D. The moderately (d)efficient enzyme: Catalysis-related damage in vivo and its repair. Biochemistry 2021, 60, 3555–3565.

(8) Bathe, U.; Leong, B. J.; Van Gelder, K.; Barbier, G. G.; Henry, C. S.; Amthor, J. S.; Hanson, A. D. Respiratory energy demands and scope for demand expansion and destruction. Plant Physiol. 2023, 191, 2093–2103.

(9) Ruszkowski, M.; Dauter, Z. Structures of Medicago truncatula L-histidinol dehydrogenase show rearrangements required for NAD+ binding and the cofactor positioned to accept a hydride. Sci. Rep. 2017, 7, 10476.

(10) Huang, Q.; Szklarczyk, D.; Wang, M.; Simonovic, M.; von Mering, C. PaxDb 5.0: Curated protein quantification data suggests adaptive proteome changes in yeasts. Mol. Cell. Proteomics 2023, 22, 100640.

(11) Adams, E. Synthesis and properties of an α-amino aldehyde, histidinal. J. Biol. Chem. 1955, 217, 317–324.

(12) Görisch, H.; Hölke, W. Binding of histidinal to histidinol dehydrogenase. Eur. J. Biochem. 1985, 150, 305–308.

(13) Nagai, A.; Kheirolomoom, A.; Ohta, D. Site-directed mutagenesis shows that the conserved cysteine residues of histidinol dehydrogenase are not essential for catalysis. J. Biochem. 1993, 114, 856–861.

(14) Ravikumar, A.; Arzumanyan, G. A.; Obadi, M. K. A.; Javanpour, A. A.; Liu, C. C. Scalable, continuous evolution of genes at mutation rates above genomic error thresholds. Cell 2018, 175, 1946–1957.

(15) Molina, R. S.; Rix, G.; Mengiste, A. A.; Álvarez, B.; Seo, D.; Chen, H.; Hurtado, J.; Zhang, Q.; García-García, J. D.; Heins, Z. J.; Almhjell, P. J.; Arnold, F. H.; Khalil, A. S.; Hanson, A. D.; Dueber, J. E.; Schaffer, D. V.; Chen, F.; Kim, S.; Fernández, L. A.; Shoulders, M. D.; Liu, C. C. In vivo hypermutation and continuous evolution with cellular systems. Nat. Rev. 2022, 2, 1–22.

(16) García-García, J. D.; Van Gelder, K.; Joshi, J.; Bathe, U.; Leong, B. J.; Bruner, S. D.; Liu, C. C.; Hanson, A. D. Using continuous directed evolution to improve enzymes for plant applications. Plant Physiol. 2022, 188, 971–983.

(17) Newton, M. S.; Arcus, V. L.; Gerth, M. L.; Patrick, W. M. Enzyme evolution: Innovation is easy, optimization is complicated. Curr. Opin. Struct. Biol. 2018, 48, 110–116.

(18) Bar-Even, A.; Noor, E.; Savir, Y.; Liebermeister, W.; Davidi, D.; Tawfik, D.S.; Milo, R. The moderately efficient enzyme: evolutionary and physicochemical trends shaping enzyme parameters. Biochemistry 2011, 50, 4402–4410.

(19) Wright, M. B.; Howell, E. A.; Gaber, R. F. Amino acid substitutions in membrane-spanning domains of Hol1, a member of the major facilitator superfamily of transporters, confer nonselective cation uptake in Saccharomyces cerevisiae. J. Bacteriol. 1996, 178, 7197–7205.

(20) Zhong, Z.; Wong, B. G.; Ravikumar, A.; Arzumanyan, G. A.; Khalil, A. S.; Liu, C. C. Automated continuous evolution of proteins in vivo. ACS Synth. Biol. 2020, 9, 1270–1276.

(21) Ravikumar, A.; Arrieta, A.; Liu, C. C. An orthogonal DNA replication system in yeast. Nat. Chem. Biol. 2014, 10, 175–177.

(22) Grubmeyer, C. T.; Insinga, S.; Bhatia, M.; Moazami, N. Salmonella typhimurium histidinol dehydrogenase: complete reaction stereochemistry and active site mapping. Biochemistry 1989, 28, 8174–8180.

(23) Kinsella, A. R.; Smith, D.; Pickard, M. Resistance to chemotherapeutic antimetabolites: a function of salvage pathway involvement and cellular response to DNA damage. Br. J. Cancer 1997, 75, 935–945.

(24) Widholm, J. Selection and characterization of amino acid analog resistant plant cell cultures. Crop Sci. 1977, 17, 597–600.

(25) Saier, M. H., Jr. Understanding the genetic code. J. Bacteriol. 2019, 201, e00091–19.

(26) Prywes, N.; Phillips, N. R.; Oltrogge, L. M.; Lindner, S.; Taylor-Kearney, L. J.; Tsai, Y. C.; de Pins, B.; Cowan, A. E.; Chang, H.A.; Wang, R. Z.; Hall, L. N.; Bellieny-Rabelo, D.; Nisonoff, H. M.; Weissman, R. F.; Flamholz, A. I.; Ding, D.; Bhatt, A. Y.; Mueller-Cajar, O.; Shih, P. M.; Milo, R.; Savage, D. F. A map of the rubisco biochemical landscape. Nature 2025, 638, 823–828.

(27) LaRue, T. A.; Spencer, J. F. The utilization of imidazoles by yeasts. Can. J. Microbiol. 1967, 13, 789–794.

(28) Han, B.; Tang, Y.; Xie, Y.; Liu, H.; Zhou, H.; Wu, S.; Zhan, J.; Huang, W.; You, Y. The characteristics of histamine and tyramine degradation of Saccharomyces cerevisiae HL10 and HL17 and their application in wine fermentation. Food Microbiol. 2025,131, 104804.

(29) Arnold, F. H. Directed evolution: bringing new chemistry to life. Angew. Chem. Int. Ed. Engl. 2018, 57, 4143–4148.

(30) Cagiada, M.; Johansson, K. E.; Valanciute, A.; Nielsen, S. V.; Hartmann-Petersen, R.; Yang, J. J.; Fowler, D. M.; Stein, A.; Lindorff-Larsen, K. Understanding the origins of loss of protein function by analyzing the effects of thousands of variants on activity and abundance. Mol. Biol. Evol. 2021, 38, 3235–3246.

(31) Denard, C. A.; Ren, H.; Zhao, H. Improving and repurposing biocatalysts via directed evolution. Curr. Opin. Chem. Biol. 2015, 25, 55–64.

(32) Van Gelder, K.; Gayen, A. K.; Hanson, A. D. Mirages in continuous directed enzyme evolution: a cautionary case study with plantized bacterial THI4 enzymes. Plant Biotechnol. J. 2025, 23, 1070–1072.

(33) Lanza, A. M.; Curran, K. A.; Rey, L. G.; Alper, H. S. A condition-specific codon optimization approach for improved heterologous gene expression in Saccharomyces cerevisiae. BMC Syst. Biol. 2014, 8, 33.

(34) Zhao, M.; Ma, J.; Zhang, L.; Qi, H. Engineering strategies for enhanced heterologous protein production by Saccharomyces cerevisiae. Microb. Cell Fact. 2024, 23, 32.

(35) Van Gelder, K.; Oliveira-Filho, E. R.; García-García, J. D.; Hu, Y.; Bruner, S. D.; Hanson, A. D. Directed evolution of aerotolerance in sulfide-dependent thiazole synthases. ACS Synth. Biol. 2023, 12, 4.

(36) Gaber, R. F.; Kielland-Brandt, M. C.; Fink, G. R. HOL1 mutations confer novel ion transport in Saccharomyces cerevisiae. Mol. Cell. Biol. 1990, 10, 643–652.

(37) García-García, J. D.; Joshi, J.; Patterson, J. A.; Trujillo-Rodriguez, L.; Reisch, C. R.; Javanpour, A. A.; Liu, C. C.; Hanson, A. D. potential for applying continuous directed evolution to plant enzymes: an exploratory study. Life 2020, 10, 179.

(38) Belt, K.; Obe, D.; Wilson, M. A.; Millar, A. H.; Bathe, U. harnessing mass spectrometry-based proteomics for continuous directed evolution. bioRxiv 2025.03.03.641153; doi: 10.1101/2025.03.03.641153.

(39) Schneider, C.; Rasband, W.; Eliceiri, K. NIH Image to ImageJ: 25 years of image analysis. Nat, Methods 2012, 9, 671–675.

(40) Tomasello, G.; Armenia, I.; Molla, G. The Protein Imager: a full-featured online molecular viewer interface with server-side HQ-rendering capabilities. Bioinformatics 2020, 36, 2909–2911.

