## Supplementary material for "Continuous Directed Evolution of a Plant Histidinol Dehydrogenase to Extend Lifespan": Proteomics Methods

**Sample Preparation for Mass Spectrometry Analysis.** Arabidopsis HDH was analyzed from both purified proteins and dried yeast extracts from evolved populations. For protein extraction, 10 mg of dried yeast was incubated in 200 µL SDS extraction buffer (7% SDS, 0.1 M Tris, 5 mM TCEP, 1× protease inhibitor, pH 7.5). Samples were heated at 95 °C for 10 min while shaking at 1000 rpm, then briefly sonicated in a water bath for 1 min to improve cell lysis. After centrifugation at maximum speed (~18,000 g) for 10 min, the supernatant was transferred to a 96-well plate for downstream processing.

**Protein Quantification and Digestion.** Protein concentrations were estimated using the bicinchoninic acid (BCA) assay. Proteins were digested following an SP3-assisted protocol,[1] using trypsin at an enzyme-to-substrate ratio of 1:50. Digestion was performed at 37 °C for 18 h to ensure thorough cleavage. A 10-µL aliquot from each digested sample was used for subsequent analysis.

**Mass Spectrometry Analysis.** For untargeted proteomic analysis, samples were injected into an online nanoflow LC-MS system operating at 0.4 µL/min using a capillary column (Picofrit, 50 µm tip, 75 µm ID; New Objective, ICT36015030F-50) packed in-house with 15 cm of C18 reverse-phase material (3 µm particle size; Dr. Maisch GmbH). The LC was coupled to a Thermo Scientific Fusion mass spectrometer via a Dionex Ultimate 3000 UHPLC system. Ionization was achieved using a spray voltage of 2 kV, with the capillary temperature maintained at 275 °C. For data-dependent acquisition (DDA), full MS scans were acquired at a resolution of 60,000 at *m/z* 200, with an AGC target of 300% and auto injection time. The MS1 scan range was set to 350–1500 *m/z*. Fragment ion spectra were acquired at a resolution of 15,000 with auto injection time, a 'standard' AGC target, and an intensity threshold of $5 \times 10^4$. The isolation width was set at 1.6 *m/z* and the normalized collision energy (NCE) was 30%. Raw MS data were processed using MaxQuant (v2.4.0.0)[2] with default settings. Trypsin was specified as the protease, allowing up to two missed cleavages. Carbamidomethylation of cysteine was set as a fixed modification, while oxidation of methionine and N-terminal acetylation were included as variable modifications. A minimum of seven peptides was required, with a maximum of five modifications. *The match between runs* feature was enabled. Label-free quantification (LFQ) was performed using the MaxQuant LFQ algorithm, with the minimum LFQ ratio count set to 2 and *fast LFQ* enabled. Protein identification was performed against a combined FASTA database containing the evolved protein sequence of *Arabidopsis* HDH and the *S. cerevisiae* reference proteome (UP000002311, UniProt). MaxQuant output files were subsequently analyzed in R.

**Development of HDH standards for targeted proteomic analysis.** To enable absolute quantification of *Arabidopsis* HDH in yeast lysates, purified *Arabidopsis* HDH was used as spike-in standard. They were expressed in *E. coli* and purified. The identity and purity of the recombinant protein was confirmed via SDS-PAGE and mass spectrometry. Known concentrations of recombinant HDH were spiked into

*S. cerevisiae* BY4742 lysates that contain no construct expressing *Arabidopsis* HDH. To ensure specificity, peptides unique to *Arabidopsis* HDH, distinct from the yeast HIS4 protein, were selected based on *in silico* digestion and peptide uniqueness analysis. These peptides were used for downstream MRM assay development and quantification. A series of spike-in concentrations were used to construct a standard curve by correlating MS signal intensities with known amounts of HDH. The standard curves enabled absolute quantification of the target proteins in experimental samples by comparing the MRM signal intensities of expressed HDH and HDH peptides to those of the spike-in standards.

**Targeted mass spectrometry analysis.** For targeted quantification via multiple reaction monitoring (MRM), digested peptides were separated on an Aeris Peptide XB-C18 column (3.6 µm, 100 Å, 50 × 2.1 mm; Phenomenex) using a Thermo UltiMate 3000 RSLCnano system coupled to a Thermo TSQ Altis triple quadrupole mass spectrometer. The column was maintained at 55 °C with a flow rate of 0.4 mL/min. The gradient elution profile was as follows:

- 0–20.5 min: 2% to 22% acetonitrile (ACN) with 0.1% formic acid (FA)
- 20.5–27.5 min: 22% to 35% ACN with 0.1% FA
- 27.5–28.5 min: 35% to 97% ACN with 0.1% FA
- 28.5–29.2 min: hold at 97% ACN with 0.1% FA
- 29.2–29.5 min: return to 2% ACN with 0.1% FA
- 29.5–35 min: re-equilibration at 2% ACN with 0.1% FA

The list of selected peptide transitions used for SRM is provided in Table S3. Targeted peak areas were quantified using Skyline (v24.1.0.199).[3] Absolute protein concentrations in the yeast samples were subsequently determined by comparing the MS signal intensities of recombinant proteins to those of the spiked-in standards. Further data analysis was performed using R.

**Supplemental References**

(1) Hughes, C. S.; Moggridge, S.; Müller, T.; Sorensen, P. H.; Morin, G  B.; Krijgsveld, J. Single-pot, solid-phase-enhanced sample preparation for proteomics experiments. *Nat Protoc*. **2019,** *14*, 68–85.

(2) Cox, J.; Mann, M. MaxQuant enables high peptide identification rates, individualized p.p.b.-range mass accuracies and proteome-wide protein quantification. *Nat Biotechnol*. **2008,** *26*, 1367–1372.

(3) MacLean, B.; Tomazela, D. M,; Shulman, N.; Chambers, M.; Finney, G. L.; Frewen, B.; Kern, R.; Tabb, D. L.; Liebler, D. C.; MacCoss, M. J. Skyline: an open source document editor for creating and analyzing targeted proteomics experiments. *Bioinformatics.* **2010,** *26*, 966–968.
