## Supplementary material for "Continuous Directed Evolution of a Plant Histidinol Dehydrogenase to Extend Lifespan": Figure S1

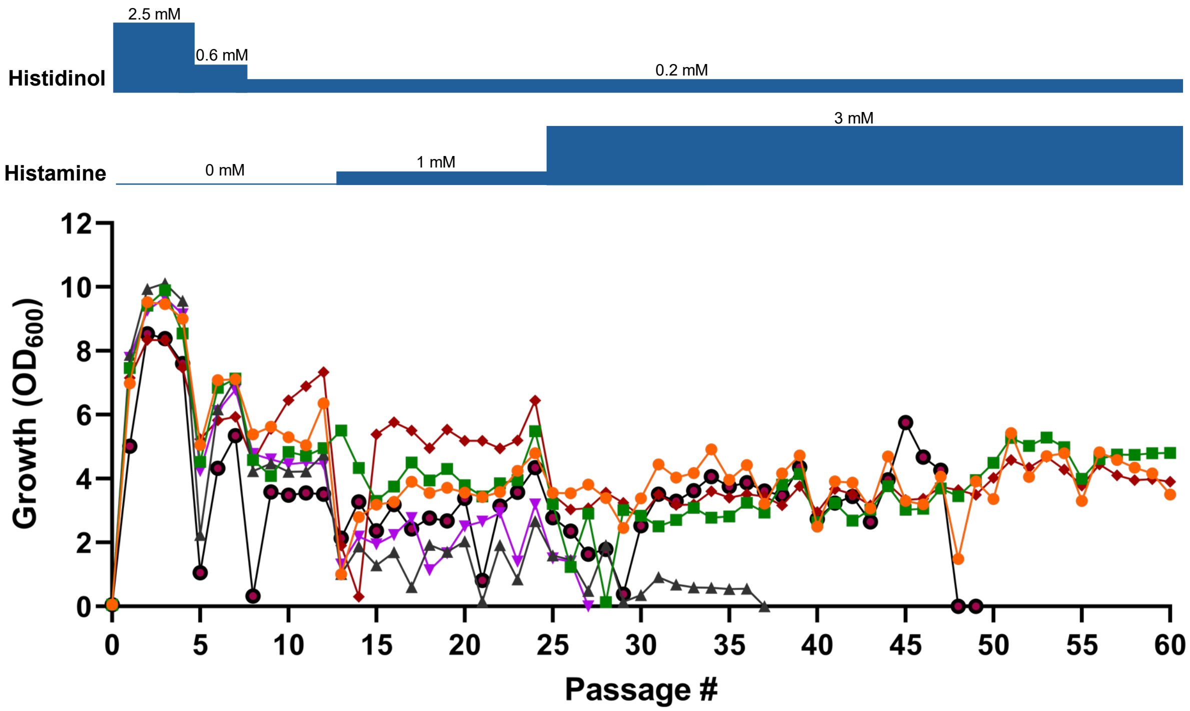


**Figure S1**. Full campaign growth profiles for representative subpopulations (in different colors) that expressed the non-polyadenylated HDH construct. Cells received stepwise decreasing levels of histidinol, and stepwise increasing levels of histamine as indicated. Passage length was generally 2-3 days but was extended to 4-5 days for the early passages in each step except the first.
