## Supplementary material for "Continuous Directed Evolution of a Plant Histidinol Dehydrogenase to Extend Lifespan": Figure S2

| | | | A72 | | | | | | | | | | | | | | | | | A0 | | | |
|---|---|---|---|---|---|---|---|---|---|---|---|---|---|---|---|---|---|---|---|---|---|---|---|
| | Position | Base change | 1 | 1.1 | 1.2 | 1.3 | 4 | 4.1 | 4.2 | 4.3 | 6 | 7 | 7.1 | 7.2 | 7.3 | 9 | 9.1 | 9.2 | 9.3 | 3 | 3.1 | 4 | 8 |
| 10B2 | N/A | N/A | | | | | | | | | | | | | | | | | | | | | |
| K2K | 3 | A>G | | | | | | | | | | | | | | | | | | | | | |
| N33D | 1 | A>G | | | | | | | | | | | | | | | | | | | | | |
| A38T | 1 | G>A | | | | | | | | | | | | | | | | | | | | | |
| R40R | 3 | A>G | | | | | | | | | | | | | | | | | | | | | |
| D75G | 2 | A>G | | | | | | | | | | | | | | | | | | | | | |
| A81A | 3 | T>C | | | | | | | | | | | | | | | | | | | | | |
| F82L | 1 | T>C | | | | | | | | | | | | | | | | | | | | | |
| D87D | 3 | C>T | | | | | | | | | | | | | | | | | | | | | |
| S98P | 1 | T>C | | | | | | | | | | | | | | | | | | | | | |
| T99T | 3 | A>G | | | | | | | | | | | | | | | | | | | | | |
| K101E | 1 | A>G | | | | | | | | | | | | | | | | | | | | | |
| S102P | 1 | T>C | | | | | | | | | | | | | | | | | | | | | |
| E104K | 1 | G>A | | | | | | | | | | | | | | | | | | | | | |
| R110R | 3 | A>G | | | | | | | | | | | | | | | | | | | | | |
| C111C | 3 | C>T | | | | | | | | | | | | | | | | | | | | | |
| R116R | 3 | A>G | | | | | | | | | | | | | | | | | | | | | |
| S120G | 1 | A>G | | | | | | | | | | | | | | | | | | | | | |
| Y124Y | 3 | C>T | | | | | | | | | | | | | | | | | | | | | |
| A130T | 1 | G>A | | | | | | | | | | | | | | | | | | | | | |
| G147G | 3 | T>C | | | | | | | | | | | | | | | | | | | | | |
| C148C | 3 | T>C | | | | | | | | | | | | | | | | | | | | | |
| I163T | 2 | T>C | | | | | | | | | | | | | | | | | | | | | |
| V176I | 1 | G>A | | | | | | | | | | | | | | | | | | | | | |
| A192A | 3 | T>C | | | | | | | | | | | | | | | | | | | | | |
| G206G | 3 | T>A | | | | | | | | | | | | | | | | | | | | | |
| I228I | 3 | T>C | | | | | | | | | | | | | | | | | | | | | |
| S235S | 3 | C>T | | | | | | | | | | | | | | | | | | | | | |
| S274P | 1 | T>C | | | | | | | | | | | | | | | | | | | | | |
| A279A | 3 | T>C | | | | | | | | | | | | | | | | | | | | | |
| E283G | 2 | A>G | | | | | | | | | | | | | | | | | | | | | |
| P292S | 1 | C>T | | | | | | | | | | | | | | | | | | | | | |
| G294D | 2 | G>A | | | | | | | | | | | | | | | | | | | | | |
| F296L | 1 | T>C | | | | | | | | | | | | | | | | | | | | | |
| K336E | 1 | A>G | | | | | | | | | | | | | | | | | | | | | |
| V367A | 2 | T>C | | | | | | | | | | | | | | | | | | | | | |
| Y371Y | 3 | C>T | | | | | | | | | | | | | | | | | | | | | |
| L386S | 2 | T>C | | | | | | | | | | | | | | | | | | | | | |
| N401D | 1 | A>G | | | | | | | | | | | | | | | | | | | | | |
| V406A | 2 | T>C | | | | | | | | | | | | | | | | | | | | | |
| I429T | 2 | T>C | | | | | | | | | | | | | | | | | | | | | |

**Figure S2.** Full listing of nonsynonymous (mid-blue) and synonymous (light-blue) mutations that swept subpopulations.
