## Supplementary material for "Continuous Directed Evolution of a Plant Histidinol Dehydrogenase to Extend Lifespan": Figure S3

| A0 Evolved populations | Mutations |
|---|---|
| A0_P60_3 | A38T, S102P, E283G |
| A0_P60_3.1 | N33D, S98P, S102P, E283G, K336E |
| A0_P60_4 | P292S |

| A72 Evolved populations | Mutations |
|---|---|
| A72_P60_1 | V176I, V406A |
| A72_P60_1.1 | E104K, V176I, V406A |
| A72_P60_1.3 | E104K, V176I, F296L, V406A |
| A72_P60_4 | D75G, S120G, A130T |
| A72_P60_4.2 | S120G, A130T |
| A72_P60_6 | V367A |
| A72_P60_7 | L386S, N401D |
| A72_P60_9.2 | K101E, I163T, S274P |

present in nature:
variable residue

not present in nature:
conserved residue
weakly conserved residue
variable residue

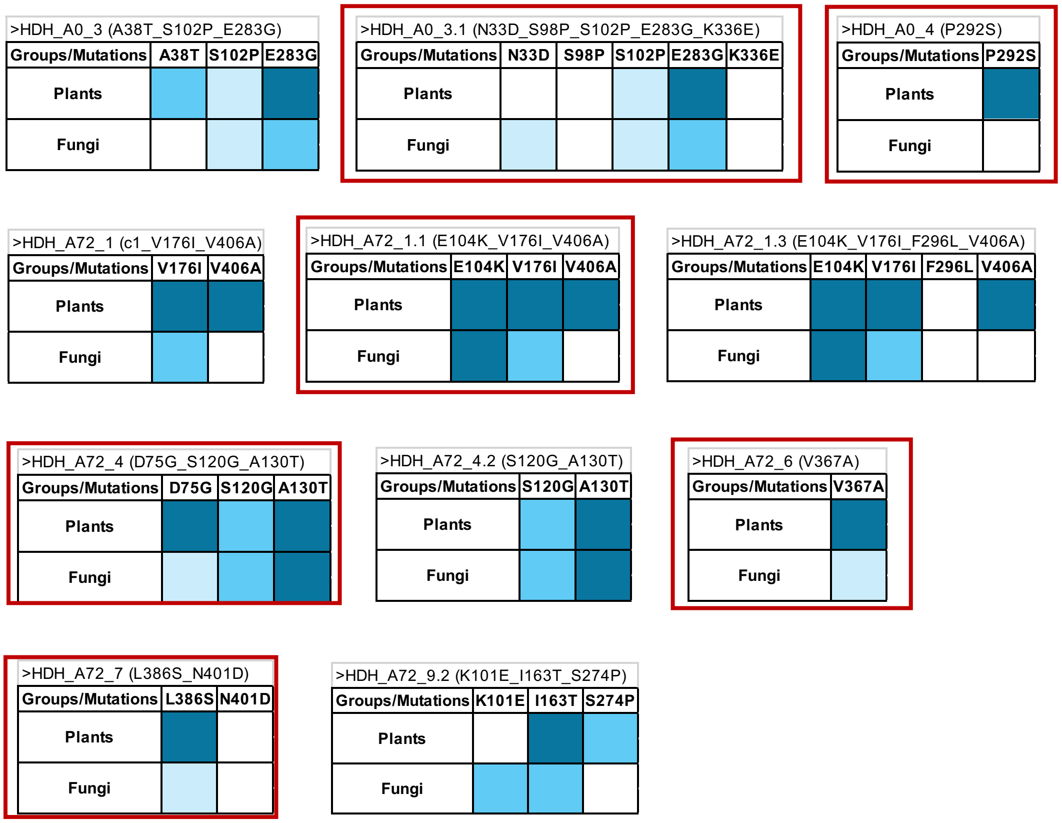


**Figure S3.** The 11 mutant HDH sequences selected for proteomics and (boxed in red) for kinetic analyses. The table lists the mutations in each sequence. The charts classify the mutations in each sequence according to their natural occurrence in plants or fungi, and the extent of natural variation at each position.
