## Supplementary material for "Continuous Directed Evolution of a Plant Histidinol Dehydrogenase to Extend Lifespan": Figure S4

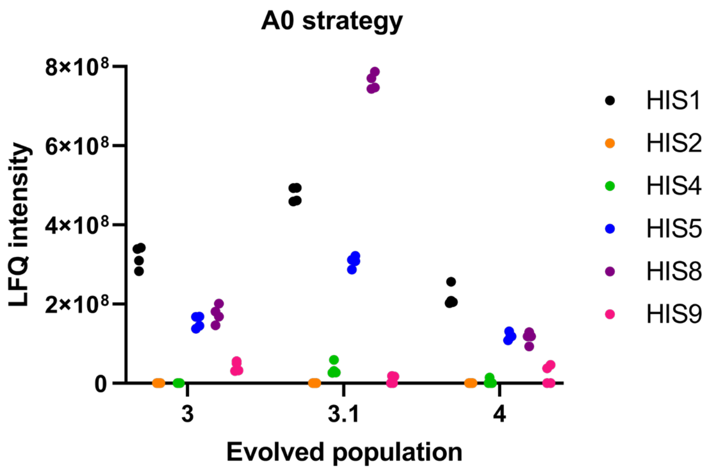


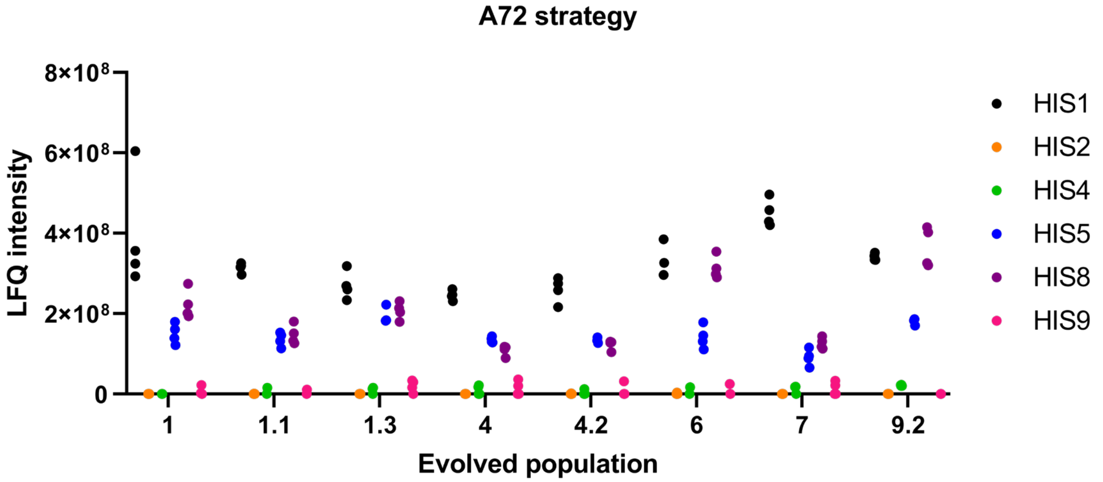


**Figure S4.** Untargeted (Orbitrap) proteomics data on levels of histidine biosynthesis enzymes in subpopulations at the end of evolution campaigns. Each symbol represents a replicate sample.
