## Supplementary material for "Continuous Directed Evolution of a Plant Histidinol Dehydrogenase to Extend Lifespan": Table S1

### Table S1. Protein and nucleotide sequences of the HDHs used in this study.

| Recoded for yeast | | |
|---|---|---|
| **Name** | **Protein sequence** | **Nucleotide sequence (5’-3’)** |
| HDH_wt<br>Recoded for yeast | MKSYRLSELSSSQVDSLKSRPRIDFSSIFATVNPIIDAVRSNGDNAVKEYTERFDKVQLNKVVEDMSELSV<br>PELDSNVKEAFDVAYDNIYAFHLAQKSTEKSVENMKGVRCKRVSRSIGSVGLYVPGGTAVLPSTALMLAIP<br>AQIAGCKTVVLATPPSKDGSICKEVLYCAKRAGVTHILKAGGAQAIAAMAWGTDSCPKVEKIFGPGNQYVT<br>AAKMILQNSEAMVSIDMPAGPSEVLVIADEHASPVYIAADLLSQAEHGPDSQVVLVVVGDSVDLNAIEEEI<br>AKQCKSLPRGEFASKALSHSFTVFARDMIEAISFSNLYAPEHLIINVKDAEKWEGLIENAGSVFIGPWTPE<br>SVGDYASGTNHVLPTYGYARMYSGVSLDSFLKFMTVQSLTEEGLRNLGPYVATMAEIEGLDAHKRAVTLRL<br>KDIEAKQLA* | ATGAAATCTTACAGATTGTCCGAATTATCTTCTTCTCAAGTCGATTCTTTGAAGTCCAGACCAAGAAT<br>CGATTTCTCGTCCATTTTTGCTACCGTTAACCCAATCATTGATGCCGTTAGATCCAACGGTGATAACG<br>CTGTCAAGGAATACACCGAAAGATTCGACAAGGTTCAACTAAACAAGGTCGTTGAAGACATGTCTGAA<br>TTGTCCGTTCCAGAATTGGACAGCAATGTCAAAGAAGCTTTTGATGTCGCTTACGACAACATCTACGC<br>TTTCCATTTAGCTCAAAAGTCCACAGAAAAGTCCGTCGAAAACATGAAGGGTGTCAGATGTAAGCGTG<br>TTTCTAGATCTATTGGTAGCGTTGGTTTGTACGTCCCAGGTGGTACCGCTGTCTTACCTTCTACTGCT<br>TTGATGTTGGCTATTCCAGCTCAAATCGCCGGTTGTAAGACCGTCGTTTTGGCTACCCCACCTTCTAA<br>GGATGGTTCTATCTGTAAGGAGGTTTTGTACTGTGCCAAGCGTGCGGGTGTTACCCACATTTTGAAGG<br>CTGGTGGTGCTCAAGCCATTGCCGCTATGGCTTGGGGTACTGACTCTTGCCCAAAGGTTGAAAAGATC<br>TTCGGTCCAGGAAACCAATACGTTACTGCTGCTAAGATGATCTTGCAAAACTCTGAAGCCATGGTTTC<br>CATTGACATGCCAGCTGGCCCATCCGAAGTCTTGGTTATCGCTGACGAACACGCTTCTCCAGTTTATA<br>TTGCTGCTGATTTGTTGTCTCAAGCCGAACACGGTCCAGACTCCCAAGTTGTCTTAGTTGTCGTGGGG<br>GATTCCGTTGACTTGAACGCTATCGAAGAAGAAATCGCTAAGCAATGTAAATCTCTGCCAAGAGGTGA<br>ATTTGCTTCCAAGGCTCTTTCCCACAGTTTCACTGTTTTCGCTAGAGACATGATTGAAGCTATCTCTT<br>TCTCTAACTTGTATGCTCCAGAACATTTGATCATCAACGTTAAGGATGCCGAAAAATGGGAAGGTTTG<br>ATTGAAAATGCTGGTTCTGTCTTCATCGGTCCATGGACTCCAGAATCTGTCGGTGACTACGCTTCCGG<br>TACCAACCACGTTTTGCCAACTTACGGTTACGCCAGAATGTACTCTGGTGTTTCTTTGGATTCCTTCT<br>TGAAGTTCATGACTGTCCAATCCTTGACTGAAGAAGGTTTGAGAAACTTGGGTCCATACGTTGCTACC<br>ATGGCTGAAATTGAAGGTCTAGACGCTCACAAGAGAGCTGTCACTTTACGTTTGAAGGACATTGAAGC<br>TAAACAATTGGCCTGA |
| Recoded for *E. coli* | | |
| **Name** | **Protein sequence** | **Nucleotide sequence (5’-3’)** |
| HDH_wt | MKSYRLSELSSSQVDSLKSRPRIDFSSIFATVNPIIDAVRSNGDNAVKEYTERFDKVQLNKVVEDMSELSV<br>PELDSNVKEAFDVAYDNIYAFHLAQKSTEKSVENMKGVRCKRVSRSIGSVGLYVPGGTAVLPSTALMLAIP<br>AQIAGCKTVVLATPPSKDGSICKEVLYCAKRAGVTHILKAGGAQAIAAMAWGTDSCPKVEKIFGPGNQYVT<br>AAKMILQNSEAMVSIDMPAGPSEVLVIADEHASPVYIAADLLSQAEHGPDSQVVLVVVGDSVDLNAIEEEI<br>AKQCKSLPRGEFASKALSHSFTVFARDMIEAISFSNLYAPEHLIINVKDAEKWEGLIENAGSVFIGPWTPE | ATGAAGTCCTATCGTCTGTCCGAGTTAAGCAGCTCCCAAGTAGACTCGCTGAAATCCAGACCCCGTAT<br>CGACTTCAGCTCGATCTTCGCCACCGTGAACCCGATTATCGACGCAGTTCGTAGCAACGGTGACAATG<br>CGGTGAAGGAGTACACCGAACGTTTTGATAAAGTGCAGCTGAATAAAGTTGTGGAAGATATGTCAGAA<br>CTGAGCGTCCCGGAGCTGGACTCGAACGTGAAGGAGGCTTTCGATGTTGCGTATGATAACATTTATGC<br>GTTTCACCTGGCGCAGAAATCGACTGAGAAGAGCGTCGAGAACATGAAAGGCGTGCGTTGCAAACGTG<br>TTAGCCGTAGCATCGGCAGCGTGGGCTTGTACGTCCCGGGTGGTACTGCCGTGCTGCCGAGCACGGCG |

| | | |
|---|---|---|
| | SVGDYASGTNHVLPTYGYARMYSGVSLDSFLKFMTVQSLTEEGLRNLGPYVATMAEIEGLDAHKRAVTLRL<br>KDIEAKQLA* | CTGATGCTGGCGATCCCGGCTCAAATTGCGGGTTGTAAGACCGTTGTTCTGGCTACCCCGCCTTCAAA<br>GGATGGTTCCATTTGCAAGGAGGTTTTATACTGCGCGAAGCGCGCAGGCGTTACCCACATTTTGAAGG<br>CTGGTGGCGCACAAGCGATTGCAGCAATGGCTTGGGGTACCGATTCTTGCCCGAAAGTTGAAAAAATC<br>TTCGGCCCGGGAAACCAGTATGTGACGGCAGCGAAGATGATCCTTCAGAATTCGGAGGCCATGGTGAG<br>TATAGATATGCCGGCGGGCCCGAGCGAAGTTTTAGTTATTGCGGACGAACACGCTTCCCCGGTCTATA<br>TCGCTGCGGATTTGCTGTCCCAAGCGGAACATGGTCCGGATTCTCAGGTTGTTCTGGTGGTGGTGGGC<br>GACTCTGTGGACCTCAATGCCATCGAGGAGGAAATTGCGAAACAGTGTAAAAGCCTACCGCGTGGTGA<br>ATTTGCGAGCAAAGCACTGAGCCATAGCTTTACCGTGTTTGCCCGCGATATGATTGAGGCGATTAGCT<br>TCTCCAACTTGTACGCGCCAGAGCACTTGATCATCAACGTTAAAGACGCGGAAAAATGGGAAGGTCTG<br>ATCGAAAACGCCGGTAGTGTGTTCATCGGTCCGTGGACCCCAGAAAGCGTGGGCGACTACGCGAGCGG<br>TACGAACCATGTCTTGCCGACCTACGGCTACGCGCGCATGTATAGCGGTGTTTCTCTGGACTCTTTTC<br>TGAAGTTCATGACGGTCCAAAGCCTGACCGAAGAGGGCCTGCGCAATCTGGGTCCGTACGTAGCCACC<br>ATGGCAGAGATCGAGGGCCTGGACGCCCATAAACGTGCGGTTACCCTCCGCTTGAAGGACATTGAGGC<br>GAAGCAGCTGGCATAA |
| HDH_A72_1.1 | MKSYRLSELSSSQVDSLKSRPRIDFSSIFATVNPIIDAVRSNGDNAVKEYTERFDKVQLNKVVEDMSELSV<br>PELDSNVKEAFDVAYDNIYAFHLAQKSTEKSVKNMKGVRCKRVSRSIGSVGLYVPGGTAVLPSTALMLAIP<br>AQIAGCKTVVLATPPSKDGSICKEVLYCAKRAGITHILKAGGAQAIAAMAWGTDSCPKVEKIFGPGNQYVT<br>AAKMILQNSEAMVSIDMPAGPSEVLVIADEHASPVYIAADLLSQAEHGPDSQVVLVVVGDSVDLNAIEEEI<br>AKQCKSLPRGEFASKALSHSFTVFARDMIEAISFSNLYAPEHLIINVKDAEKWEGLIENAGSVFIGPWTPE<br>SVGDYASGTNHVLPTYGYARMYSGVSLDSFLKFMTVQSLTEEGLRNLGPYAATMAEIEGLDAHKRAVTLRL<br>KDIEAKQLA* | ATGAAGTCCTATCGTCTGTCCGAGTTAAGCAGCTCCCAAGTAGACTCGCTGAAATCCAGACCCCGTAT<br>CGACTTCAGCTCGATCTTCGCCACCGTGAACCCGATTATCGACGCAGTTCGTAGCAACGGTGACAATG<br>CGGTGAAGGAGTACACCGAACGTTTTGATAAAGTGCAGCTGAATAAAGTTGTGGAAGATATGTCAGAA<br>CTGAGCGTCCCGGAGCTGGACTCGAACGTGAAGGAGGCTTTCGATGTTGCGTATGATAACATTTATGC<br>GTTTCACCTGGCGCAGAAATCGACTGAGAAGAGCGTCAAAAACATGAAAGGCGTGCGTTGCAAACGTG<br>TTAGCCGTAGCATCGGCAGCGTGGGCTTGTACGTCCCGGGTGGTACTGCCGTGCTGCCGAGCACGGCG<br>CTGATGCTGGCGATCCCGGCTCAAATTGCGGGTTGTAAGACCGTTGTTCTGGCTACCCCGCCTTCAAA<br>GGATGGTTCCATTTGCAAGGAGGTTTTATACTGCGCGAAGCGCGCAGGCATTACCCACATTTTGAAGG<br>CTGGTGGCGCACAAGCGATTGCAGCAATGGCTTGGGGTACCGATTCTTGCCCGAAAGTTGAAAAAATC<br>TTCGGCCCGGGAAACCAGTATGTGACGGCAGCGAAGATGATCCTTCAGAATTCGGAGGCCATGGTGAG<br>TATAGATATGCCGGCGGGCCCGAGCGAAGTTTTAGTTATTGCGGACGAACACGCTTCCCCGGTCTATA<br>TCGCTGCGGATTTGCTGTCCCAAGCGGAACATGGTCCGGATTCTCAGGTTGTTCTGGTGGTGGTGGGC<br>GACTCTGTGGACCTCAATGCCATCGAGGAGGAAATTGCGAAACAGTGTAAAAGCCTACCGCGTGGTGA<br>ATTTGCGAGCAAAGCACTGAGCCATAGCTTTACCGTGTTTGCCCGCGATATGATTGAGGCGATTAGCT<br>TCTCCAACTTGTACGCGCCAGAGCACTTGATCATCAACGTTAAAGACGCGGAAAAATGGGAAGGTCTG<br>ATCGAAAACGCCGGTAGTGTGTTCATCGGTCCGTGGACCCCAGAAAGCGTGGGCGACTACGCGAGCGG<br>TACGAACCATGTCTTGCCGACCTACGGCTACGCGCGCATGTATAGCGGTGTTTCTCTGGACTCTTTTC<br>TGAAGTTCATGACGGTCCAAAGCCTGACCGAAGAGGGCCTGCGCAATCTGGGTCCGTACGCAGCCACC |

| | | |
|---|---|---|
| | | ATGGCAGAGATCGAGGGCCTGGACGCCCATAAACGTGCGGTTACCCTCCGCTTGAAGGACATTGAGGC<br>GAAGCAGCTGGCATAA |
| HDH_A72_4 | MKSYRLSELSSSQVDSLKSRPRIDFSSIFATVNPIIDAVRSNGDNAVKEYTERFDKVQLNKVVEDMSELSV<br>PELGSNVKEAFDVAYDNIYAFHLAQKSTEKSVENMKGVRCKRVSRSIGGVGLYVPGGTTVLPSTALMLAIP<br>AQIAGCKTVVLATPPSKDGSICKEVLYCAKRAGVTHILKAGGAQAIAAMAWGTDSCPKVEKIFGPGNQYVT<br>AAKMILQNSEAMVSIDMPAGPSEVLVIADEHASPVYIAADLLSQAEHGPDSQVVLVVVGDSVDLNAIEEEI<br>AKQCKSLPRGEFASKALSHSFTVFARDMIEAISFSNLYAPEHLIINVKDAEKWEGLIENAGSVFIGPWTPE<br>SVGDYASGTNHVLPTYGYARMYSGVSLDSFLKFMTVQSLTEEGLRNLGPYVATMAEIEGLDAHKRAVTLRL<br>KDIEAKQLA* | ATGAAGTCCTATCGTCTGTCCGAGTTAAGCAGCTCCCAAGTAGACTCGCTGAAATCCAGACCCCGTAT<br>CGACTTCAGCTCGATCTTCGCCACCGTGAACCCGATTATCGACGCAGTTCGTAGCAACGGTGACAATG<br>CGGTGAAGGAGTACACCGAACGTTTTGATAAAGTGCAGCTGAATAAAGTTGTGGAAGATATGTCAGAA<br>CTGAGCGTCCCGGAGCTGGGCTCGAACGTGAAGGAGGCTTTCGATGTTGCGTATGATAACATTTATGC<br>GTTTCACCTGGCGCAGAAATCGACTGAGAAGAGCGTCGAGAACATGAAAGGCGTGCGTTGCAAACGTG<br>TTAGCCGTAGCATCGGCGGCGTGGGCTTGTACGTCCCGGGTGGTACTACCGTGCTGCCGAGCACGGCG<br>CTGATGCTGGCGATCCCGGCTCAAATTGCGGGTTGTAAGACCGTTGTTCTGGCTACCCCGCCTTCAAA<br>GGATGGTTCCATTTGCAAGGAGGTTTTATACTGCGCGAAGCGCGCAGGCGTTACCCACATTTTGAAGG<br>CTGGTGGCGCACAAGCGATTGCAGCAATGGCTTGGGGTACCGATTCTTGCCCGAAAGTTGAAAAAATC<br>TTCGGCCCGGGAAACCAGTATGTGACGGCAGCGAAGATGATCCTTCAGAATTCGGAGGCCATGGTGAG<br>TATAGATATGCCGGCGGGCCCGAGCGAAGTTTTAGTTATTGCGGACGAACACGCTTCCCCGGTCTATA<br>TCGCTGCGGATTTGCTGTCCCAAGCGGAACATGGTCCGGATTCTCAGGTTGTTCTGGTGGTGGTGGGC<br>GACTCTGTGGACCTCAATGCCATCGAGGAGGAAATTGCGAAACAGTGTAAAAGCCTACCGCGTGGTGA<br>ATTTGCGAGCAAAGCACTGAGCCATAGCTTTACCGTGTTTGCCCGCGATATGATTGAGGCGATTAGCT<br>TCTCCAACTTGTACGCGCCAGAGCACTTGATCATCAACGTTAAAGACGCGGAAAAATGGGAAGGTCTG<br>ATCGAAAACGCCGGTAGTGTGTTCATCGGTCCGTGGACCCCAGAAAGCGTGGGCGACTACGCGAGCGG<br>TACGAACCATGTCTTGCCGACCTACGGCTACGCGCGCATGTATAGCGGTGTTTCTCTGGACTCTTTTC<br>TGAAGTTCATGACGGTCCAAAGCCTGACCGAAGAGGGCCTGCGCAATCTGGGTCCGTACGTAGCCACC<br>ATGGCAGAGATCGAGGGCCTGGACGCCCATAAACGTGCGGTTACCCTCCGCTTGAAGGACATTGAGGC<br>GAAGCAGCTGGCATAA |
| HDH_A72_6 | MKSYRLSELSSSQVDSLKSRPRIDFSSIFATVNPIIDAVRSNGDNAVKEYTERFDKVQLNKVVEDMSELSV<br>PELDSNVKEAFDVAYDNIYAFHLAQKSTEKSVENMKGVRCKRVSRSIGSVGLYVPGGTAVLPSTALMLAIP<br>AQIAGCKTVVLATPPSKDGSICKEVLYCAKRAGVTHILKAGGAQAIAAMAWGTDSCPKVEKIFGPGNQYVT<br>AAKMILQNSEAMVSIDMPAGPSEVLVIADEHASPVYIAADLLSQAEHGPDSQVVLVVVGDSVDLNAIEEEI<br>AKQCKSLPRGEFASKALSHSFTVFARDMIEAISFSNLYAPEHLIINVKDAEKWEGLIENAGSVFIGPWTPE<br>SVGDYASGTNHALPTYGYARMYSGVSLDSFLKFMTVQSLTEEGLRNLGPYVATMAEIEGLDAHKRAVTLRL<br>KDIEAKQLA* | ATGAAGTCCTATCGTCTGTCCGAGTTAAGCAGCTCCCAAGTAGACTCGCTGAAATCCAGACCCCGTAT<br>CGACTTCAGCTCGATCTTCGCCACCGTGAACCCGATTATCGACGCAGTTCGTAGCAACGGTGACAATG<br>CGGTGAAGGAGTACACCGAACGTTTTGATAAAGTGCAGCTGAATAAAGTTGTGGAAGATATGTCAGAA<br>CTGAGCGTCCCGGAGCTGGACTCGAACGTGAAGGAGGCTTTCGATGTTGCGTATGATAACATTTATGC<br>GTTTCACCTGGCGCAGAAATCGACTGAGAAGAGCGTCGAGAACATGAAAGGCGTGCGTTGCAAACGTG<br>TTAGCCGTAGCATCGGCAGCGTGGGCTTGTACGTCCCGGGTGGTACTGCCGTGCTGCCGAGCACGGCG<br>CTGATGCTGGCGATCCCGGCTCAAATTGCGGGTTGTAAGACCGTTGTTCTGGCTACCCCGCCTTCAAA<br>GGATGGTTCCATTTGCAAGGAGGTTTTATACTGCGCGAAGCGCGCAGGCGTTACCCACATTTTGAAGG<br>CTGGTGGCGCACAAGCGATTGCAGCAATGGCTTGGGGTACCGATTCTTGCCCGAAAGTTGAAAAAATC<br>TTCGGCCCGGGAAACCAGTATGTGACGGCAGCGAAGATGATCCTTCAGAATTCGGAGGCCATGGTGAG<br>TATAGATATGCCGGCGGGCCCGAGCGAAGTTTTAGTTATTGCGGACGAACACGCTTCCCCGGTCTATA |

| | | |
|---|---|---|
| | | TCGCTGCGGATTTGCTGTCCCAAGCGGAACATGGTCCGGATTCTCAGGTTGTTCTGGTGGTGGGC<br>GACTCTGTGGACCTCAATGCCATCGAGGAGGAAATTGCGAAACAGTGTAAAAGCCTACCGCGTGGTGA<br>ATTTGCGAGCAAAGCACTGAGCCATAGCTTTACCGTGTTTGCCCGCGATATGATTGAGGCGATTAGCT<br>TCTCCAACTTGTACGCGCCAGAGCACTTGATCATCAACGTTAAAGACGCGGAAAAATGGGAAGGTCTG<br>ATCGAAAACGCCGGTAGTGTGTTCATCGGTCCGTGGACCCCAGAAAGCGTGGGCGACTACGCGAGCGG<br>TACGAACCATGCCTTGCCGACCTACGGCTACGCGCGCATGTATAGCGGTGTTTCTCTGGACTCTTTTC<br>TGAAGTTCATGACGGTCCAAAGCCTGACCGAAGAGGGCCTGCGCAATCTGGGTCCGTACGTAGCCACC<br>ATGGCAGAGATCGAGGGCCTGGACGCCCATAAACGTGCGGTTACCCTCCGCTTGAAGGACATTGAGGC<br>GAAGCAGCTGGCATAA |
| HDH_A72_7 | MKSYRLSELSSSQVDSLKSRPRIDFSSIFATVNPIIDAVRSNGDNAVKEYTERFDKVQLNKVVEDMSELSV<br>PELDSNVKEAFDVAYDNIYAFHLAQKSTEKSVENMKGVRCKRVSRSIGSVGLYVPGGTAVLPSTALMLAIP<br>AQIAGCKTVVLATPPSKDGSICKEVLYCAKRAGVTHILKAGGAQAIAAMAWGTDSCPKVEKIFGPGNQYVT<br>AAKMILQNSEAMVSIDMPAGPSEVLVIADEHASPVYIAADLLSQAEHGPDSQVVLVVVGDSVDLNAIEEEI<br>AKQCKSLPRGEFASKALSHSFTVFARDMIEAISFSNLYAPEHLIINVKDAEKWEGLIENAGSVFIGPWTPE<br>SVGDYASGTNHVLPTYGYARMYSGVSLDSFSKFMTVQSLTEEGLRDLGPYVATMAEIEGLDAHKRAVTLRL<br>KDIEAKQLA* | ATGAAGTCCTATCGTCTGTCCGAGTTAAGCAGCTCCCAAGTAGACTCGCTGAAATCCAGACCCCGTAT<br>CGACTTCAGCTCGATCTTCGCCACCGTGAACCCGATTATCGACGCAGTTCGTAGCAACGGTGACAATG<br>CGGTGAAGGAGTACACCGAACGTTTTGATAAAGTGCAGCTGAATAAAGTTGTGGAAGATATGTCAGAA<br>CTGAGCGTCCCGGAGCTGGACTCGAACGTGAAGGAGGCTTTCGATGTTGCGTATGATAACATTTATGC<br>GTTTCACCTGGCGCAGAAATCGACTGAGAAGAGCGTCGAGAACATGAAAGGCGTGCGTTGCAAACGTG<br>TTAGCCGTAGCATCGGCAGCGTGGGCTTGTACGTCCCGGGTGGTACTGCCGTGCTGCCGAGCACGGCG<br>CTGATGCTGGCGATCCCGGCTCAAATTGCGGGTTGTAAGACCGTTGTTCTGGCTACCCCGCCTTCAAA<br>GGATGGTTCCATTTGCAAGGAGGTTTTATACTGCGCGAAGCGCGCAGGCGTTACCCACATTTTGAAGG<br>CTGGTGGCGCACAAGCGATTGCAGCAATGGCTTGGGGTACCGATTCTTGCCCGAAAGTTGAAAAAATC<br>TTCGGCCCGGGAAACCAGTATGTGACGGCAGCGAAGATGATCCTTCAGAATTCGGAGGCCATGGTGAG<br>TATAGATATGCCGGCGGGCCCGAGCGAAGTTTTAGTTATTGCGGACGAACACGCTTCCCCGGTCTATA<br>TCGCTGCGGATTTGCTGTCCCAAGCGGAACATGGTCCGGATTCTCAGGTTGTTCTGGTGGTGGGC<br>GACTCTGTGGACCTCAATGCCATCGAGGAGGAAATTGCGAAACAGTGTAAAAGCCTACCGCGTGGTGA<br>ATTTGCGAGCAAAGCACTGAGCCATAGCTTTACCGTGTTTGCCCGCGATATGATTGAGGCGATTAGCT<br>TCTCCAACTTGTACGCGCCAGAGCACTTGATCATCAACGTTAAAGACGCGGAAAAATGGGAAGGTCTG<br>ATCGAAAACGCCGGTAGTGTGTTCATCGGTCCGTGGACCCCAGAAAGCGTGGGCGACTACGCGAGCGG<br>TACGAACCATGTCTTGCCGACCTACGGCTACGCGCGCATGTATAGCGGTGTTTCTCTGGACTCTTTTT<br>CGAAGTTCATGACGGTCCAAAGCCTGACCGAAGAGGGCCTGCGCGATCTGGGTCCGTACGTAGCCACC<br>ATGGCAGAGATCGAGGGCCTGGACGCCCATAAACGTGCGGTTACCCTCCGCTTGAAGGACATTGAGGC<br>GAAGCAGCTGGCATAA |
| HDH_A0_3.1 | MKSYRLSELSSSQVDSLKSRPRIDFSSIFATVDPIIDAVRSNGDNAVKEYTERFDKVQLNKVVEDMSELSV<br>PELDSNVKEAFDVAYDNIYAFHLAQKPTEKPVENMKGVRCKRVSRSIGSVGLYVPGGTAVLPSTALMLAIP<br>AQIAGCKTVVLATPPSKDGSICKEVLYCAKRAGVTHILKAGGAQAIAAMAWGTDSCPKVEKIFGPGNQYVT<br>AAKMILQNSEAMVSIDMPAGPSEVLVIADEHASPVYIAADLLSQAEHGPDSQVVLVVVGDSVDLNAIEEGI | ATGAAGTCCTATCGTCTGTCCGAGTTAAGCAGCTCCCAAGTAGACTCGCTGAAATCCAGACCCCGTAT<br>CGACTTCAGCTCGATCTTCGCCACCGTGGACCCGATTATCGACGCAGTTCGTAGCAACGGTGACAATG<br>CGGTGAAGGAGTACACCGAACGTTTTGATAAAGTGCAGCTGAATAAAGTTGTGGAAGATATGTCAGAA<br>CTGAGCGTCCCGGAGCTGGACTCGAACGTGAAGGAGGCTTTCGATGTTGCGTATGATAACATTTATGC |

| | | |
|---|---|---|
| | AKQCKSLPRGEFASKALSHSFTVFARDMIEAISFSNLYAPEHLIINVKDAEEWEGLIENAGSVFIGPWTPE<br>SVGDYASGTNHVLPTYGYARMYSGVSLDSFLKFMTVQSLTEEGLRNLGPYVATMAEIEGLDAHKRAVTLRL<br>KDIEAKQLA* | GTTTCACCTGGCGCAGAAACCGACTGAGAAGCCCGTCGAGAACATGAAAGGCGTGCGTTGCAAACGTG<br>TTAGCCGTAGCATCGGCAGCGTGGGCTTGTACGTCCCGGGTGGTACTGCCGTGCTGCCGAGCACGGCG<br>CTGATGCTGGCGATCCCGGCTCAAATTGCGGGTTGTAAGACCGTTGTTCTGGCTACCCCGCCTTCAAA<br>GGATGGTTCCATTTGCAAGGAGGTTTTATACTGCGCGAAGCGCGCAGGCGTTACCCACATTTTGAAGG<br>CTGGTGGCGCACAAGCGATTGCAGCAATGGCTTGGGGTACCGATTCTTGCCCGAAAGTTGAAAAAATC<br>TTCGGCCCGGGAAACCAGTATGTGACGGCAGCGAAGATGATCCTTCAGAATTCGGAGGCCATGGTGAG<br>TATAGATATGCCGGCGGGCCCGAGCGAAGTTTTAGTTATTGCGGACGAACACGCTTCCCCGGTCTATA<br>TCGCTGCGGATTTGCTGTCCCAAGCGGAACATGGTCCGGATTCTCAGGTTGTTCTGGTGGTGGTGGGC<br>GACTCTGTGGACCTCAATGCCATCGAGGAGGGAATTGCGAAACAGTGTAAAAGCCTACCGCGTGGTGA<br>ATTTGCGAGCAAAGCACTGAGCCATAGCTTTACCGTGTTTGCCCGCGATATGATTGAGGCGATTAGCT<br>TCTCCAACTTGTACGCGCCAGAGCACTTGATCATCAACGTTAAAGACGCGGAAGAATGGGAAGGTCTG<br>ATCGAAAACGCCGGTAGTGTGTTCATCGGTCCGTGGACCCCAGAAAGCGTGGGCGACTACGCGAGCGG<br>TACGAACCATGTCTTGCCGACCTACGGCTACGCGCGCATGTATAGCGGTGTTTCTCTGGACTCTTTTC<br>TGAAGTTCATGACGGTCCAAAGCCTGACCGAAGAGGGCCTGCGCAATCTGGGTCCGTACGTAGCCACC<br>ATGGCAGAGATCGAGGGCCTGGACGCCCATAAACGTGCGGTTACCCTCCGCTTGAAGGACATTGAGGC<br>GAAGCAGCTGGCATAA |
| HDH_A0_4 | MKSYRLSELSSSQVDSLKSRPRIDFSSIFATVNPIIDAVRSNGDNAVKEYTERFDKVQLNKVVEDMSELSV<br>PELDSNVKEAFDVAYDNIYAFHLAQKSTEKSVENMKGVRCKRVSRSIGSVGLYVPGGTAVLPSTALMLAIP<br>AQIAGCKTVVLATPPSKDGSICKEVLYCAKRAGVTHILKAGGAQAIAAMAWGTDSCPKVEKIFGPGNQYVT<br>AAKMILQNSEAMVSIDMPAGPSEVLVIADEHASPVYIAADLLSQAEHGPDSQVVLVVVGDSVDLNAIEEEI<br>AKQCKSLSRGEFASKALSHSFTVFARDMIEAISFSNLYAPEHLIINVKDAEKWEGLIENAGSVFIGPWTPE<br>SVGDYASGTNHVLPTYGYARMYSGVSLDSFLKFMTVQSLTEEGLRNLGPYVATMAEIEGLDAHKRAVTLRL<br>KDIEAKQLA* | ATGAAGTCCTATCGTCTGTCCGAGTTAAGCAGCTCCCAAGTAGACTCGCTGAAATCCAGACCCCGTAT<br>CGACTTCAGCTCGATCTTCGCCACCGTGAACCCGATTATCGACGCAGTTCGTAGCAACGGTGACAATG<br>CGGTGAAGGAGTACACCGAACGTTTTGATAAAGTGCAGCTGAATAAAGTTGTGGAAGATATGTCAGAA<br>CTGAGCGTCCCGGAGCTGGACTCGAACGTGAAGGAGGCTTTCGATGTTGCGTATGATAACATTTATGC<br>GTTTCACCTGGCGCAGAAATCGACTGAGAAGAGCGTCGAGAACATGAAAGGCGTGCGTTGCAAACGTG<br>TTAGCCGTAGCATCGGCAGCGTGGGCTTGTACGTCCCGGGTGGTACTGCCGTGCTGCCGAGCACGGCG<br>CTGATGCTGGCGATCCCGGCTCAAATTGCGGGTTGTAAGACCGTTGTTCTGGCTACCCCGCCTTCAAA<br>GGATGGTTCCATTTGCAAGGAGGTTTTATACTGCGCGAAGCGCGCAGGCGTTACCCACATTTTGAAGG<br>CTGGTGGCGCACAAGCGATTGCAGCAATGGCTTGGGGTACCGATTCTTGCCCGAAAGTTGAAAAAATC<br>TTCGGCCCGGGAAACCAGTATGTGACGGCAGCGAAGATGATCCTTCAGAATTCGGAGGCCATGGTGAG<br>TATAGATATGCCGGCGGGCCCGAGCGAAGTTTTAGTTATTGCGGACGAACACGCTTCCCCGGTCTATA<br>TCGCTGCGGATTTGCTGTCCCAAGCGGAACATGGTCCGGATTCTCAGGTTGTTCTGGTGGTGGTGGGC<br>GACTCTGTGGACCTCAATGCCATCGAGGAGGAAATTGCGAAACAGTGTAAAAGCCTATCGCGTGGTGA<br>ATTTGCGAGCAAAGCACTGAGCCATAGCTTTACCGTGTTTGCCCGCGATATGATTGAGGCGATTAGCT<br>TCTCCAACTTGTACGCGCCAGAGCACTTGATCATCAACGTTAAAGACGCGGAAAAATGGGAAGGTCTG<br>ATCGAAAACGCCGGTAGTGTGTTCATCGGTCCGTGGACCCCAGAAAGCGTGGGCGACTACGCGAGCGG<br>TACGAACCATGTCTTGCCGACCTACGGCTACGCGCGCATGTATAGCGGTGTTTCTCTGGACTCTTTTC |

| | | |
|---|---|---|
| | | TGAAGTTCATGACGGTCCAAAGCCTGACCGAAGAGGGCCTGCGCAATCTGGGTCCGTACGTAGCCACC<br>ATGGCAGAGATCGAGGGCCTGGACGCCCATAAACGTGCGGTTACCCTCCGCTTGAAGGACATTGAGGC<br>GAAGCAGCTGGCATAA |
