## Supplementary material for "Continuous Directed Evolution of a Plant Histidinol Dehydrogenase to Extend Lifespan": Table S2

**Table S2. Primers used in this study.**

| Primer name | Sequence (5'-3') | Purpose |
|---|---|---|
| AtHDH_*Nde*I_FW | CAGTCACATATGAAGTCCTATCGTCTGTCCG | Cloning of HDH into pET28b(+) |
| AtHDH_*Hind*III_RV | AGTCTGAAGCTTTTATGCCAGCTGCTTCGCCTC | Cloning of HDH into pET28b(+) |
| HA_LB_FW | GGCTACGGTCTCGGGAGGGAGCTTTCGCCTAGCGAG | Cloning of HOL1 deletion cassette |
| HA_LB_RV | GGCTACGGTCTCAAACTGTTAAAAATTTAAGATTCACACTCGC | Cloning of HOL1 deletion cassette |
| promoter_FW | GGCTACGGTCTCAAGTTTAAGAGCTTGGTGAGCG | Cloning of HOL1 deletion cassette |
| promoter_RV | GGCTACGGTCTCTTTTGCCTTCGTTTATCTTGC | Cloning of HOL1 deletion cassette |
| LEU2_FW | GGCTACGGTCTCTCAAAGATGCACGTTCCTAAGAAGATC | Cloning of HOL1 deletion cassette |
| LEU2_RV | GGCTACGGTCTCCAAGCAAGGATTTTCTTAACTTCTTC | Cloning of HOL1 deletion cassette |
| terminator_FW | GGCTACGGTCTCCGCTTAACACCGATTATTTAAAGCTGCTG | Cloning of HOL1 deletion cassette |
| terminator_RV | GGCTACGGTCTCCACAAGAAAGACGTCTCTGTATGATCCGTCGAGTTCAAG | Cloning of HOL1 deletion cassette |
| HA_RB_FW | GGCTACGGTCTCCTTGTATCTAGTAGATGATAGGAATTAAG | Cloning of HOL1 deletion cassette |
| HA_RB_RV | GGCTACGGTCTCCAGCGCCAGAATCTCAGAAGGGAG | Cloning of HOL1 deletion cassette |
| seqpAGT572_FW | CCAATCTAAGTCTGTGCTC | Sequencing HOL1 deletion cassette |
| seqpAGT572_RV | CAGCTTATCATCGATAACGG | Sequencing HOL1 deletion cassette |
| promSEQ_FW | CCAGGTATCGTTTGAACACG | Sequencing HOL1 deletion cassette |
| promSEQ_RV | TAGAGTGTACTAGAGGAGGC | Sequencing HOL1 deletion cassette |
| LEU2seq1_FW | GTACGCCAACTTAAGACC | Sequencing HOL1 deletion cassette |
| LEU2seq_RV | ATGGTGGCTCATGTTGTAGG | Sequencing HOL1 deletion cassette |
| LEU2seq2_FW | TGGATGCAGGTATCAGAACTGG | Sequencing HOL1 deletion cassette |
| termSEQ_RV | GCAGCAGCTTTAAATAATCGG | Sequencing HOL1 deletion cassette |
| HOL1delcst_FW | GGAGCTTTCGCCTAGCGAG | Sequencing HOL1 deletion cassette |
| HOL1delcst_RV | CCAGAATCTCAGAAGGGAG | Sequencing HOL1 deletion cassette |
| HOL1_HA1_FW | GGAGCTTTCGCCTAGCGAGAAC | Cloning of HOL1 into pGEM; Producing HOL1-V509F cassette |

| | | |
|---|---|---|
| HOL1_HA2_RV | CCAGAATCTCAGAAGGGAGAAACC | Cloning of HOL1 into pGEM; Producing HOL1-V509F cassette |
| HOL1_V509F_F | GATTGTTACCCCGATATGTTTTTGGAAGGTATGG | SDM for HOL1-1 mutation (V509F) |
| HOL1_V509F_R | CATTAAATACGCCATCGCAATATCACC | SDM for HOL1-1 mutation (V509F) |
| HOL1_ext_F | CATCAAGGCCTGGTTGTGTC | Amplifying HOL1 for sequencing |
| HOL1_ext_R | GTTTGTCTATCGTATCGCAGCC | Amplifing HOL1 for sequencing |
| HOL1_seq1F | GCTATTACTACCAAGTTGACATCC | Sequencing HOL1 |
| HOL1_seq2F | GATCGCTGGGTACATATCTGC | Sequencing HOL1 |
| HOL1_seq3F | GAATGTTCCCACACTTATTGGAGC | Sequencing HOL1 |
| HOL1_seq1R | CCACGTGTTTCACGAAGGGC | Sequencing HOL1 |
| HOL1_seq2R | CATGGAGGTTCGTAGTAGGC | Sequencing HOL1 |
| HOL1_seq3R | GTCAATACGGAGCCCAATTGGTGC | Sequencing HOL1 |
| p1_new_FW | GAGATTATTGGAAGATTAGTACGTCTCC | Amplifying HDH from p1 |
| Linker_LEU_RV | CCTCCCACTAATTCTCTGACAACAACG | Amplifying HDH from p1 |
| 10B2_FW | GAGATTATTGGAAGATTAGTACGTCTCC | Sequencing HDH in p1 |
| LEU_short_RV | CCTCCCACTAATTCTCTGACAACAACG | Sequencing HDH in p1 |
| HDH.Sc_intF | CCAATACGTTACTGCTGC | Sequencing HDH in p1 |
| HDH.Sc_int2F | ATGCTGGTTCTGTCTTCATCG | Sequencing HDH in p1 |
| HDH.Sc_intR | GAGAAGCGTGTTCGTCAGC | Sequencing HDH in p1 |
| HDH_L386S_FW | GGACTCTTTTagcAAGTTCATGACG | SDM for HDH_A72_7 |
| HDH_L386S_RV | AGAGAAACACCGCTATAC | SDM for HDH_A72_7 |
| HDH_N401D_FW | GGGCCTGCGCgatCTGGGTCCGT | SDM for HDH_A72_7 |
| HDH_N401D_RV | TCTTCGGTCAGGCTTTGGACCGTC | SDM for HDH_A72_7 |
| HDH_P292S_FW | TAAAAGCCTATCGCGTGGTGAATTTGC | SDM for HDH_A0_4 |
| HDH_P292S_RV | CACTGTTTCGCAATTTC | SDM for HDH_A0_4 |
| ScHDH_cds_FW | ATGGTTTTGCCGATTCTACCG | Verifying *his4* knock-out in yeast |

| | | |
|---|---|---|
| ScHDH_cds_RV | CTACTGGAAATCCTTTGGGATCAACC | Verifying *his4* knock-out in yeast |
| ext_ScHDH_FW | ATATCATAGCACAACTGCGCTGTG | Verifying *his4* knock-out in yeast |
| ext_ScHDH_RV | CATTCATTACATAGTGTATCTCTATTCATTCAAGAC | Verifying *his4* knock-out in yeast |
