## Supplementary material for "Continuous Directed Evolution of a Plant Histidinol Dehydrogenase to Extend Lifespan": Table S3

**Table S3. Peptide sequences and mass spectrometry analysis parameters used for targeted selected reaction monitoring analysis of Arabidopsis HDH.**

| PeptideA3A3:G27 | Precursor Mz | Precursor Charge | Product Mz | Product Charge | Fragment Ion | Retention Time |
|---|---|---|---|---|---|---|
| LSELSSSQVDSLK | 696.864471 | 2 | 1063.563 | 1 | y10 | 11.48 |
| LSELSSSQVDSLK | 696.864471 | 2 | 950.47892 | 1 | y9 | 11.48 |
| LSELSSSQVDSLK | 696.864471 | 2 | 863.44689 | 1 | y8 | 11.49 |
| LSELSSSQVDSLK | 696.864471 | 2 | 776.41486 | 1 | y7 | 11.48 |
| LSELSSSQVDSLK | 696.864471 | 2 | 330.16596 | 1 | b3 | 11.48 |
| VVEDMSELSVPELDSNVK | 995.487895 | 2 | 1416.7217 | 1 | y13 | 18.24 |
| VVEDMSELSVPELDSNVK | 995.487895 | 2 | 1200.647 | 1 | y11 | 18.24 |
| VVEDMSELSVPELDSNVK | 995.487895 | 2 | 1087.563 | 1 | y10 | 18.26 |
| VVEDMSELSVPELDSNVK | 995.487895 | 2 | 1000.531 | 1 | y9 | 18.27 |
| VVEDMSELSVPELDSNVK | 995.487895 | 2 | 901.46254 | 1 | y8 | 18.26 |
| SVENMK | 354.173264 | 2 | 620.30722 | 1 | y5 | 1.95 |
| SVENMK | 354.173264 | 2 | 521.23881 | 1 | y4 | 1.95 |
| SVENMK | 354.173264 | 2 | 430.19324 | 1 | b4 | 1.84 |
| SVENMK | 354.173264 | 2 | 561.23372 | 1 | b5 | 1.95 |
| TVVLATPPSK | 506.805499 | 2 | 812.48763 | 1 | y8 | 9.15 |
| TVVLATPPSK | 506.805499 | 2 | 713.41922 | 1 | y7 | 9.15 |
| TVVLATPPSK | 506.805499 | 2 | 600.33515 | 1 | y6 | 9.15 |
| TVVLATPPSK | 506.805499 | 2 | 428.25036 | 1 | y4 | 9.15 |
| TVVLATPPSK | 506.805499 | 2 | 201.12337 | 1 | b2 | 9.15 |
| IFGPGNQYVTAAK | 683.361701 | 2 | 1105.5636 | 1 | y11 | 12.31 |
| IFGPGNQYVTAAK | 683.361701 | 2 | 1048.5422 | 1 | y10 | 12.31 |
| IFGPGNQYVTAAK | 683.361701 | 2 | 390.23471 | 1 | y4 | 12.31 |
| IFGPGNQYVTAAK | 683.361701 | 2 | 553.28546 | 2 | y11 | 12.31 |
| IFGPGNQYVTAAK | 683.361701 | 2 | 524.77473 | 2 | y10 | 12.32 |
| MYSGVSLDSFLK | 673.83667 | 2 | 1215.6256 | 1 | y11 | 20.56 |
| MYSGVSLDSFLK | 673.83667 | 2 | 1052.5623 | 1 | y10 | 20.58 |
| MYSGVSLDSFLK | 673.83667 | 2 | 965.53022 | 1 | y9 | 20.58 |
| MYSGVSLDSFLK | 673.83667 | 2 | 809.44035 | 1 | y7 | 20.56 |
| MYSGVSLDSFLK | 673.83667 | 2 | 295.11109 | 1 | b2 | 20.58 |
| FMTVQSLTEEGLR | 755.882141 | 2 | 1232.6481 | 1 | y11 | 18.31 |
| FMTVQSLTEEGLR | 755.882141 | 2 | 1131.6004 | 1 | y10 | 18.31 |
| FMTVQSLTEEGLR | 755.882141 | 2 | 1032.532 | 1 | y9 | 18.31 |
| FMTVQSLTEEGLR | 755.882141 | 2 | 904.47344 | 1 | y8 | 18.31 |
| FMTVQSLTEEGLR | 755.882141 | 2 | 704.35734 | 1 | y6 | 18.31 |
